# Colon cancer arises from differentiated cell lineages in the context of inflammation

**DOI:** 10.1101/2023.10.02.560432

**Authors:** Mathijs P. Verhagen, Rosalie Joosten, Mark Schmitt, Niko Välimäki, Andrea Sacchetti, Kristiina Rajamäki, Jiahn Choi, Paola Procopio, Sara Silva, Berdine van der Steen, Thierry P.P. van den Bosch, Danielle Seinstra, Michail Doukas, Leonard H. Augenlicht, Lauri A. Aaltonen, Riccardo Fodde

## Abstract

According to conventional views, colon cancer originates from stem cells. However, inflammation, a key risk factor for colon cancer, was shown to suppress intestinal stemness. Here, we employed Paneth cells (PCs) as a model to assess the capacity of differentiated lineages to trigger tumorigenesis in the context of inflammation. Upon inflammation, PC-specific *Apc* mutations led to intestinal tumors reminiscent not only of those arising in inflammatory bowel disease (IBD) patients but also of a larger fraction of sporadic colon cancers. The latter is likely due to the inflammatory consequences of Western-style dietary habits, the major colon cancer risk factor. Computational methods designed to predict the cell-of-origin of cancer confirmed that, in a substantial fraction of sporadic colon cancers the cells-of-origin are secretory lineages and not stem cells.

**One-Sentence Summary:** Secretory cell lineages trigger tumor formation in the context of the major etiologic colon cancer risk factors.

## Paneth cells as the origin of intestinal tumors in the context of inflammation and loss of *Apc*

The origin of the vast majority of cancers is thought to reside in stem-or progenitor-like cells which satisfy the needs for active proliferation, self-renewal, and differentiation capacity. This was demonstrated in the intestine where loss-of-function mutations in the *Apc* tumor suppressor gene successfully initiate adenoma formation only when they occur in *Lgr5*^+^ CBCs. When the same *Apc* mutation is introduced in more committed and shorter-lived transit-amplifying cells, tumor formation is halted at early micro-adenoma stages(*1*). However, next to this “bottom-up” scenario, additional “top-down” models of intestinal tumorigenesis have been proposed where committed intestinal cells initiate cancer especially in the context of tissue injury and inflammation(*2, 3*). In the specific case of colon cancer, western style dietary habits and chronic inflammation, two of the major etiologic factors associated with increased risk for sporadic malignant disease in the digestive tract, are thought to induce specific cellular and molecular alterations of the intestinal epithelium which ultimately lead to the expansion of cell targets for tumor initiation and progression(*2, 4*). However, which specific cell lineages are capable of dedifferentiating upon tissue injury and which are the underlying mechanisms remain largely unclear. Paneth cells are specialized secretory cells located at the very bottom of the crypt of Lieberkühn in the small intestine where they secrete antimicrobial peptides into the lumen(*5*). Moreover, they provide essential physical support and secrete signals to ensure *Lgr5*^+^ stem cell function(*6*). In the colon, where Paneth cells are not present, *cKit*^+^ Paneth-like cells, also known as deep crypt secretory (DCS) cells, play secretory and niche-like functional roles analogous to those of *bona fide* PCs in the small intestine(*7, 8*).

We first reported on the ability of Paneth cells to re-enter the cell cycle and de-differentiate upon irradiation and inflammation to acquire stem cell-like features and contribute to the tissue regenerative response(*9–11*). Consequently, we here questioned whether Paneth cells could be the origin of intestinal cancer upon inflammation. To this aim, we bred mice carrying Lox-alleles at genes most frequently mutated along the adenoma-to-carcinoma sequence, namely *Apc*(*12*)*, Kras*(*13*), and *Trp53*(*14*), each combined with a Cre specific for *Lgr5*^+^ CBCs (*Lgr5*^CreERT2-EGFP^)(*15*) or for Paneth cells (*Lyz1*^CreERT2^)(*16*). Following Cre activation by tamoxifen, DSS was administered through the drinking water to model inflammation (Fig. 1a). In the absence of DSS-induced inflammation, PC-specific single gene mutations did not give rise to intestinal tumors. In contrast, loss of *Apc* in *Lgr5*^+^-CBCs transformed crypts into β-catenin^hi^ foci that grew into multiple adenomas 4-6 weeks after Cre induction (Fig. 1b). When single gene mutations were combined with DSS administration, *Apc* loss in Paneth cells resulted in increased nuclear and cytoplasmic β-catenin expression eventually leading to the formation of PC-derived adenomas (Fig. 1c). Of note, Paneth-specific *Kras* or *Trp53* mutations did not result in tumor formation even in the presence of the inflammatory stimulus (Fig. 1d). However, the compound loss of *Apc* and oncogenic activation of *Kras* in Paneth cells resulted in a striking increase in tumor multiplicity (6.1 fold) even in the absence of DSS (6.9 fold)(Fig. 1e). The combination of *Apc* and *Tp53* mutations in PCs also led to an increase in tumor multiplicity upon DSS administration (1.6 fold), though to a lesser extent when compared with the compound *Apc*/*Kras*-mutant genotype, possibly indicating a distinct mechanism underlying tumor onset in these mice. Indeed, phospho-histone H2A.X (Ser 139) IHC analysis confirmed an increase in DNA damage and chromosomal instability in the *Tp53*-mutant tumors (Suppl. Fig. 1a). Targeting all three genes in Paneth cells resulted in a very aggressive phenotype with high tumor multiplicity (10.1 fold) in the absence of the inflammatory stimulus (Fig. 1e). When compared to *Apc*-driven tumors originated in PCs, the histology of adenomas from mice in which two or three genes were targeted revealed a progressive increase in dysplasia and invasive morphology (Suppl. Fig. 1b). Distribution of adenomas along the small intestine also followed distinct patterns with a prevalence of duodenal tumors in compound *Apc*/*Kras* tumors regardless of DSS (Suppl. Fig. 2a).

**Figure 1.**
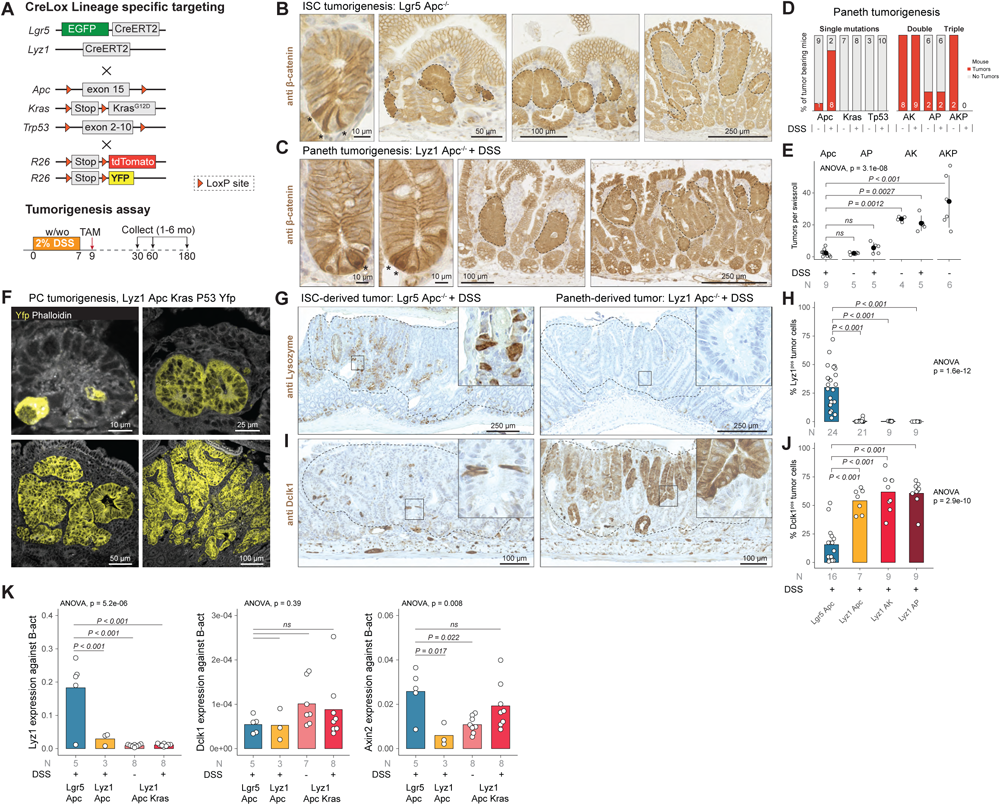
Paneth cells as the cell-of-origin of intestinal cancer. **a.** Cre-Lox strategy taimed at the targeting of *Apc*, *Kras*, and *Tp53* mutations in intestinal stem cells (*Lgr5*^+^ ISCs) and Paneth cells (*Lyz1*^+^ PCs). **b.** and **c.** β-catenin IHC analysis of intestinal tumors initiated from *Lgr5*^+^ ISCs (**b**) and PCs (**c**). The asterisks indicate *Lgr5*^+^ ISCs and PCs with enhanced cytoplasmic and nuclear β- catenin accumulation; tumor foci and adenomas are indicated by dashed lines. **d-e.** Tumor multiplicity was calculated according to tumor-bearing animals (**d**) and by tumor number per genotype (**e**) in the presence/absence of DSS and based on Swiss roll counts. Error bars denote standard deviations of the mean. P values denote one-way ANOVA and Tukey post-hoc tests for group comparisons. **f.** Lineage tracing analysis of Paneth cells (labelled by YFP) at different stages of tumor-initiation and progression. **g-j.** Left panels: Lyz1 (**g**) and Dclk1 (**i**) IHC analysis *Lgr5*^+^ ISCs and PCs-derived adenomas. Top-right inlets (**h-j**): quantification of number of *Lyz1*-(**h**) and *Dclk1*-positive (**j**) tumor cells. P values depict one-way ANOVA and Tukey post-hoc tests for group comparisons. **k.** *Lyz1*, *Dclk1*, and *Axin2* qPCR expression analysis across different adenoma genotypes. P values represent one-way ANOVA and Tukey post-hoc tests for group comparisons.

To validate the Paneth cell origin of the observed intestinal tumors, we bred *Lyz1*^CreERT2^ mice with R26^LSL-tdTomato^ or R26^LSL-YFP^ reporters and traced their lineage upon tamoxifen-driven targeting of the *Apc, Kras,* and *Tp53* mutations. As shown in Figure 1f and Suppl. Fig. 2b, this confirmed the Paneth cell origin of the corresponding tumors by capturing the process from microscopic lesions to adenoma formation.

Overall, these results demonstrate that Paneth cells can initiate intestinal adenomas upon genetic ablation of *Apc* in the context of inflammation. In combination with *Apc* loss, activation of oncogenic *Kras* or loss of *Tp53* function rescues the need for an inflammatory stimulus and results in increased PC-derived tumor multiplicities and progression to malignant phenotype.

Next, we characterized lineage-specific markers in the PC-and CBC-originated tumors by immunohistochemistry. Of note, while cells expressing the PC marker lysozyme (Lyz1) were notable in *Lgr5*-derived tumors (*Lgr5*/*Apc*: 30.0 % ± 18.5 positive tumor cells), they were nearly absent in adenomas that originated from Paneth cells (*Lyz1*/*Apc*: 0.48 % ± 1.16)(Fig. 1g-h). The opposite was observed for Dclk1 (doublecortin like kinase 1), a Tuft(*17*) and tumor stem cell marker(*18, 19*), that was more frequently detected among PC-derived adenomas (*Lyz1*/*Apc*: 54.1 % ± 10.5) when compared with *Lgr5*-derived tumors (*Lgr5*/*Apc*: 15.6 % ± 15.7)(Fig. 1i-j). Other lineage-specific markers for entero-endocrine (Chga), goblet (Muc2), and stem cells (Olfm4) showed variable levels without clear-cut differences among tumors with different cells-of-origin (Suppl. Fig. 2c). The increased *Dclk1* expression in PC-derived tumors is of interest in view of its association with increased immune and stromal infiltration in colon cancer(*20*).

To confirm these results at the transcriptional level, expression of the *Lyz1* and *Dclk1* genes was analyzed by RTqPCR (Fig. 1k). Indeed, *Lyz1* expression was lower in Paneth-derived tumors (*Lgr5*/*Apc* vs. *Lyz1*/*Apc*: log2FC = 2.64, Pval = 7.5e-4) when compared to *Lgr5*-derived tumors. *Dclk1* expression was very low and variable at the RNA level, and did not show significant differences across the groups.

To assess the relative activation levels of the Wnt signaling pathway among the different tumor groups, we measured expression levels of *Axin2*, a well-established Wnt downstream target. *Axin2* expression was higher in *Lgr5*-compared to PC-derived tumors (*Lgr5*/*Apc* vs. *Lyz1*/*Apc*: log2FC = 2.12, Pval = 0.017)(Fig. 1k). Moreover, both *Kras* oncogenic activation and inflammation gradually increased *Axin2* levels in PC-derived tumors, in agreement with the previously reported synergism between *Apc* and *Kras* mutations in the activation of the Wnt pathway(*21*).

Thus, upon tumorigenesis, Paneth cells de-differentiate to a state that hampers secretory differentiation leading to specific patterns of tumor histology and gene expression distinct from that of *Lgr5*-derived tumors.

### Paneth cells dedifferentiate into revival stem cells upon enhanced Wnt signaling activation

To elucidate the mechanisms which underlie the conversion of PCs into cells of origin of small intestinal tumors in the context of inflammation and/or of specific genetic hits, we combined scRNAseq analysis with lineage tracing. To this aim, we induced the *Apc*, *Kras*, and *Tp53* genetic mutations in the R26^LSL-tdTomato^/*Lyz1*^CreERT2^ (or R26^LSL-YFP^) reporter strains in the presence or absence of DSS (Fig. 2a). Subsequently, cells were harvested from the intestinal epithelium, purified by FACS, and transcriptionally profiled by scRNAseq (Methods; Suppl. Fig. 3). After preprocessing, we obtained the transcriptomes of 23.231 epithelial cells from 32 mice, distributed over the different lineages of the intestinal epithelium (Fig. 2b). Close examination of the cells positive for the reporter genes revealed novel clusters of PCs that arise upon DSS administration and the specific gene mutations, but were not observed among Paneth cells in homeostatic conditions (PC cluster 1-4, Fig. 2c).

**Figure 2.**
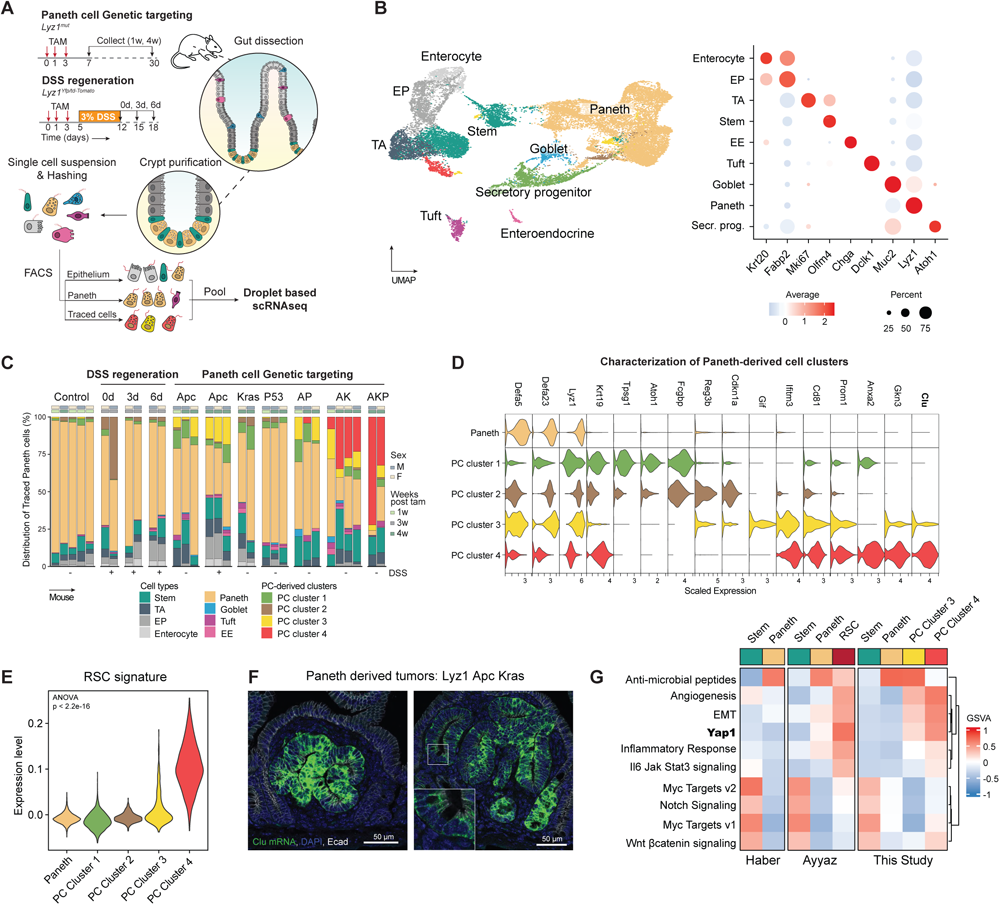
Paneth cells dedifferentiate into a revival stem cell (RSC) identity. **a.** Schematics of the experimental approach. After genetic targeting of Paneth cells, intestinal crypts were extracted, and the isolated cells labeled with hashing antibodies and sorted according to three different sorting strategies: epithelium, PC-enriched and PC-traced cells. **b.** UMAP embedding of the different cell clusters/lineages (left panel), annotated according to the expression of canonical marker genes (right panel). **c.** Bar plot of the distribution of traced cells across the different mouse genotypes and experimental conditions. **d.** Violin plots representing marker genes of the newly identified Paneth-derived cell clusters (PC Cluster 1-4). **e.** Association analysis of the revival stem cell (RSC) signature with PC Cluster 1-4. **f.** RNA *in situ* hybridizations of the *Clu* gene in small tumors derived from Paneth cells upon compound targeting of *Apc* and *Kras* mutations. **g.** Gene sets variation analysis among the Haber et al.(*72*), Ayyaz et al.(*28*), and the present studies.

To characterize the novel PC-derived states, we performed differential expression analysis and identified cluster-specific markers (Fig. 2d)(Suppl. Table 1). While PC Cluster 1 appeared at low frequency across different genotypes, PC Cluster 2 arises directly upon exposure to the inflammatory stimulus. Both PC Clusters 1 and 2 are characterized by increased expression of two markers of radio-resistant and secretory progenitors with self-renewal capacity during regeneration, namely *Krt19*(*22*) and *Atoh1*(*23*), while increased expression of *Reg3b*, known for its protective role in the development of colitis and ileitis(*24*), and *Cdkn1a* (p21), a marker of terminal differentiation in the intestine(*25*), was observed in Cluster 2 compared to 1.

PC Cluster 3 became apparent in mice carrying *Apc* mutations (7.23% ± 5.77 of traced cells) alone and in combination with DSS treatment (16.40% ± 2.10 of traced cells), and in double and triple mutant (AP, AK, AKP) animals, though not in mice carrying single *Kras* or *Tp53* mutations. PC Cluster 3 showed increased expression of *Gif* (gastric intrinsic factor), *Cd81*, a tetraspanin family member known to mark the response to γ−irradiation and correlated with the expression of ISC-and proliferation genes(*26*), and *Prom1* (prominin 1, also known as Cd133), a well-established colon cancer stem cell marker(*27*).

PC Cluster 4 consisted of cells from mice where double (AK) and triple (AKP) mutations were targeted to Paneth cells (23.25% ± 11.23 and 52.26% ± 28.23 of traced cells, respectively). Increased expression of *Anxa2* (annexin 2), a functional marker of inflammatory response, and *Clu* (clusterin), previously shown to earmark revival stem cells (RSCs) upon γ-irradiation(*28*), feature PC Cluster 4. Accordingly, evaluation of the RSC signature showed elevated expression among the PC clusters (Fig. 2e), and *in situ* hybridization analysis in PC-derived tumors from mice carrying compound mutations (AK and AKP)(Fig. 2f and Suppl. Fig. 4a-b) confirmed the increased *Clu* expression. Finally, pathway analysis revealed the similarities between the PC-derived Cluster 4 and RSCs, both earmarked by the activation of Yap1 signaling and specific inflammatory pathways (Fig. 2g). Compared to RSCs, PC-derived and *Apc/Kras-*mutant cells from Cluster 4 showed increased levels of Tgf-β and Wnt signaling (Suppl. Fig. 4c).

Thus, upon genetic targeting or inflammatory stimulus, PCs escape their homeostatic identity and acquire distinct cellular features as shown by scRNAseq and FACS analysis (Suppl. Fig. 4d-e). Of note, DSS treatment led to a lower expression of *Lgr5* and *Ascl2* in stem cells, as well as a lower association of the CBC signature, confirming ours and others’ previous observations according to which resident stem cells lose their multipotency upon acute inflammation (Suppl. Fig. 4f)(*11*).

Collectively, these findings demonstrate that PCs efficiently dedifferentiate upon genetic targeting or inflammatory stimulus leading to distinct cellular identities. During tumorigenesis driven by *Apc/Kras*, PCs share features with the Yap1-dependent revival stem cell identity, and further activate Tgf-β and Wnt signaling in their conversion to bona fide tumor cells.

### Transcriptomic comparison of Paneth-and Lgr5-derived tumors reveals a dichotomy in stem cell phenotypes

To investigate the consequences of the cell-of-origin identity on the transcriptional profile of the resulting intestinal tumors, we performed bulk RNA sequencing of macroscopically dissected lesions originated from ISCs and PCs (Fig. 3a). Principal component analysis revealed that the major variance component (61%) was attributed to differences in the cell-of-origin, while the impact of genotype or inflammatory stimulus became notable in the second component of variation (10%)(Fig. 3b). Differential expression analysis between tumors derived from PCs and ISCs in the same genetic and inflammatory context (*Apc* and DSS) revealed tumor signatures specific for each cell-of-origin (Suppl. Table 2).

**Figure 3.**
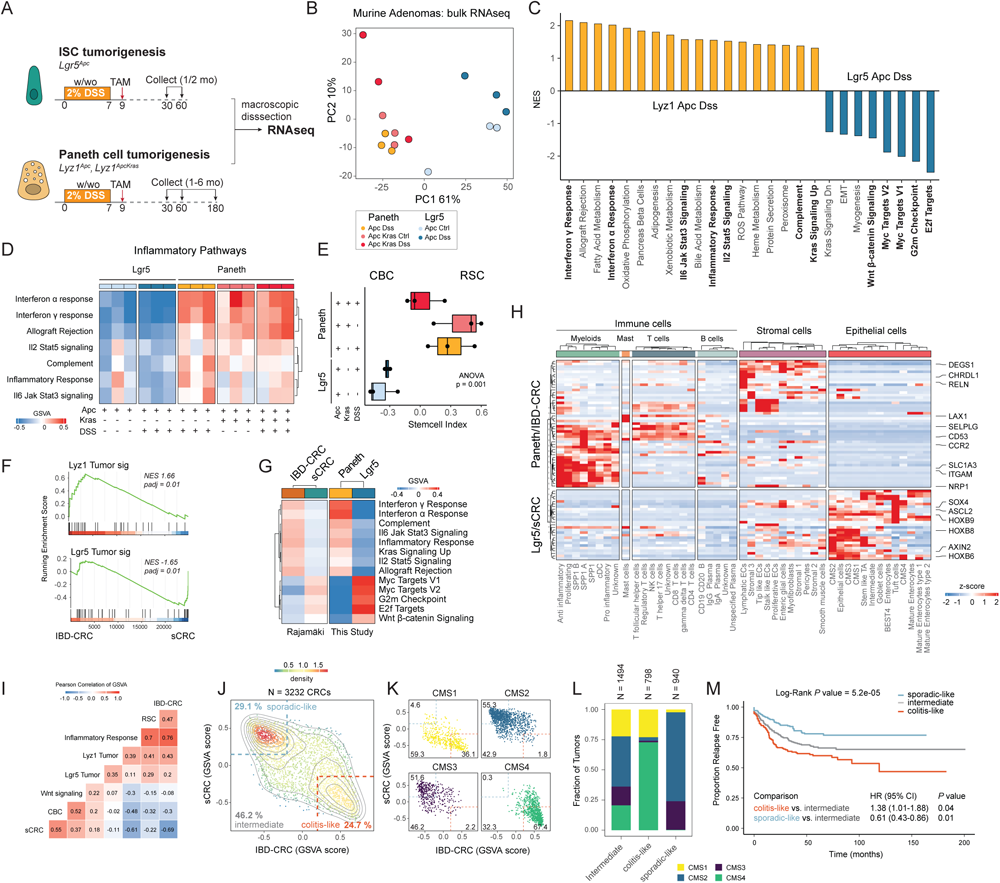
Paneth-derived adenomas have an inflammatory phenotype mimicking colitis-associated colon cancer. **a.** Schematics of the experimental approach to compare PC-and *Lgr5*-derived adenomas. **b**. Principal Component Analysis (PCA) plot showing that the cell-of-origin is the dominant discriminator of variance. **c.** Bar plot summarizing the gene set enrichment analysis between Paneth-(*Lyz1 Apc* DSS) and ISC-(*Lgr5 Apc* DSS) derived tumors. Pathways were filtered based on Pval < 0.05 and abs NES > 0.5. **d**. Subset of inflammatory pathways, visualized as heat map based on values from the gene set variation analysis. **e**. Box plots showing results of the stem cell index. P value depicts result of one-way ANOVA. **f**. Gene set enrichment analysis showing significant but opposite association between the *Lyz1* tumor signature and IBD-CRCs, and between the *Lgr5* tumor signature and sCRC. **g**. Heatmap showing GSVA scores, averaged per tumor group, of pathways with similar patterns between the murine and human tumor groups. **h**. Heatmap highlighting the differentially expressed genes (log2FC >1.5, Padj < 0.01) shared between the Paneth/IBD-tumors and the *Lgr5*/sCRC-tumors. Values denote z-scores of average expression per cell type. **i-m**. Two distinct sporadic colon cancer identities become apparent upon analysis of a large cohort of CRC tumors (N = 3232 samples). **i.** Heatmap showing Pearson Correlations of the GSVA scores. **j-k**. Scatter plot showing two distinct clusters of sporadic-and colitis-like in all (**j**) colon cancers and (**k**) stratified according to their consensus molecular subtype (CMS). Grey lines indicate contours lines, dashed lines show thresholds to classify tumors in colitis-like, sporadic-like, and intermediate groups. **l**. Stacked bar plot analysis showing the distribution of consensus molecular subtypes (CMS1 to 4) across the colitis-like and sporadic-like colon cancers. **m**. Kaplan-Meier survival analysis for relapse free survival. P values denote result of log-rank test and cox regression models for univariate analyses. Hazard ratios (HR) and confidence intervals (CI) are displayed for pairwise comparisons.

Gene set enrichment analysis (GSEA) indicated that tumors derived from ISCs were characterized by high levels of Myc and Wnt signaling, while PC-derived adenomas showed higher levels of inflammatory pathways indicative of infiltration from the tumor micro-environment (Fig. 3c, Suppl. Fig. 5). Of note, the inflammatory characteristics of PC-derived tumors were observed also in mice where *Apc* and *Kras* mutations were targeted to Paneth cells in the absence of DSS-driven inflammation (Fig. 3d), indicating that specific mutant genotypes and the type of cell-of-origin can trigger tumor initiation by mimicking the inflammatory context otherwise brought about by DSS.

Next, we employed the ISC index(*29*) (intestinal <u>s</u>tem <u>c</u>ell index; see Methods) to predict the relative proportions of RSC and CBC stem cells in the intestinal tumors. In agreement with the scRNAseq analysis, PC-derived tumors were RSC-enriched whereas the ISC-derived tumors consisted mainly of CBCs (Fig. 3e). Notably, the highest RSC contribution was observed in tumors originated from *Apc/Kras-*mutant PCs in the absence of inflammation, when compared with the equivalent genotype upon DSS administration. The latter is of relevance to dissect the relative contribution of the inflammatory insult and of the somatic mutations in the de-differentiation process leading to tumor initiation. As mentioned before, one of the primary effects of various forms of tissue injury to the intestinal epithelium is the loss of resident stem cells. In the Singh et al. (2022) study(*30*), ablation of *Lgr5*^+^ ISCs was performed to study its consequences by scRNAseq. Our analysis of these data sets revealed elevated expression of secretory genes in the regenerated stem cells normally restricted to the Paneth cell lineage (Suppl. Fig. 6a). In parallel, upon *Lgr5*^+^ cell ablation, Paneth and goblet cells partially activate the RSC program (Suppl. Fig. 6b), indicating that stem cell loss is sufficient to trigger plasticity and RSC reprogramming from PCs. Hence, in order to assess whether the removal of the resident stem cells is sufficient to activate the lineage tracing capacity of Paneth cells in the absence of the inflammatory injury, we implemented a similar DTR-based *Lgr5*^+^ ablation experiments. As shown in Suppl. Fig. 6c-d, lineage tracing from Paneth cells was observed similar to what previously shown upon inflammation(*11*).

Together, these results indicate that the cell-of-origin embodies the major source of inter-tumor variability, and that the Paneth-or ISC-origin is reflected by the RSC-or CBC-like profile of the resulting tumors, respectively. Notably, the inflammation-driven ISC loss activates de-differentiation at the epithelial level.

### The transcriptional profile of Paneth-derived tumors mimics colitis-associated colorectal cancer

The small intestinal location of Paneth cells and of the tumors originating from them raise questions on the relevance of our study for colon cancer. To explore the generalizability of our results, we first analyzed bulk and single-cell RNAseq data from two distinct studies(*31, 32*) centered around the AOM/DSS mouse model of colitis-driven colon cancer(*33*). This protocol relies on the oncogenic β-catenin mutations caused by azoxymethane (AOM) which, in combination with the DSS-driven ulcerative colitis, result in multiple adenocarcinomas in the distal colon. As shown in Suppl. Fig. 7a (by CiberSortx(*34*); see Methods), based on their bulk RNAseq data(*31*) AOM/DSS-derived colon tumors encompass an abundant population similar to the RSC-like PC Cluster 4. Moreover, analysis of the scRNAseq data(*32*) confirmed that AOM/DSS-derived colon tumors, when compared to their sporadic counterparts, were distinct in terms of the qualitative and quantitative composition of their tumor microenvironment, namely a pronounced presence of infiltrating immune cells and tumor-associated fibroblasts (Suppl. Fig. 7b). Accordingly, tumor cells derived in the context of AOM/DSS share transcriptional similarity with Paneth-derived tumor cells and RSCs (Suppl. Fig. 7c-d).

Hence, notwithstanding the small intestinal location of the PC-derived tumors, their gene expression signatures and overall inflammatory TME profiles are characteristic of colitis-associated carcinoma in the mouse.

Next, in view of the marked differences in the transcriptional profiles between mouse intestinal tumors with distinct cells of origin, we questioned whether similar differences distinguish human sporadic colon cancers from those that arose in the context of IBD. To this aim, RNAseq profiles of human microsatellite stable sporadic colon cancers (sCRC, N = 38) and from patients suffering from inflammatory bowel disease (IBD-CRC, N = 14) were interrogated(*35*). GSEA of the most differentially expressed genes (log2FC > 5, Padj < 0.01) from the mouse tumors (*Lyz*T signature, N = 27 genes; *Lgr5*T signature, N = 40 genes) revealed a significant association between the *Lyz*T signature and IBD-CRC (NES 1.67, Padj 6.2e-3), while the *Lgr5*T profile was significantly associated with sCRC (NES -1.62, Padj 0.013)(Fig. 3f, Suppl. Fig. 8a-c, Suppl. Table 3). Evaluation of the hallmarks from the molecular signature database(*36*) revealed gene sets common to PC-derived tumors and IBD-related CRCs (Interferon α/γ, inflammatory response, Il6/Il2 signaling, Kras, Complement, Allograft rejection), and to *Lgr5*-derived tumors and sporadic CRCs (Myc targets, G2m Checkpoint, E2F targets and Wnt β-catenin signaling)(Fig. 3g, Suppl. Fig. 8d).

We then compared the differentially expressed genes from PC-versus *Lgr5*-derived mouse tumors and human IBD-CRC versus sCRC (Paneth/IBD-CRC, N = 49 genes, Lgr5/sCRC: N = 27 genes) and visualized their expression across different cell types based on a large scRNAseq CRC study(*37*) (Fig. 3h). Of note, the markers shared between PC-derived tumors and IBD-CRC were dominantly expressed in myofibroblasts (e.g. *ITGAM*, *SLC1A3*), T cells (e.g. *SELPLG, LAX1*), and stromal cells (e.g. *CHRDL1, RELN*). In contrast, *Lgr5*- derived tumors/sCRC markers were mostly observed in epithelial cells (e.g. *HOXB6, HOXB8, HOXB9, AXIN2, ASCL2*), indicating the difference in stromal composition among these tumors (Fig. 3h). Comparison of a set of gene signatures (Suppl. Table 4) in a large colon cancer cohort(*38*) confirmed the presence of two distinct identities (Fig. 3i-j): a colitis-like identity enriched with RSCs and prevalent in CMS4 (67%) and CMS1 (36%) tumors, and a sporadic-like identity enriched with CBCs and common in CMS2 (55%) and CMS3 tumors (52%)(Fig. 3k-l, Suppl. Fig. 8e). Survival analysis revealed significant differences in relapse free survival between the sporadic-and colitis-like CRC groups (Pvalue = 5.2e-05)(Fig. 3m).

Thus, transcriptional signatures derived from small intestinal mouse tumors originated from Paneth cells significantly overlap with those from human colon cancer that arose in the context of IBD, possibly revealing a common cell-of-origin in secretory lineages.

### Western-style diet triggers an inflammatory response in intestinal secretory cells leading to their cell cycle activation and dedifferentiation

The relative representation of patient-derived colon cancers whose expression profiles are reminiscent of the PC-derived mouse tumors (∼25%; Figure 3j) vastly exceeds the expected proportion of colon cancers arising in patients with an history of clinically manifest IBD (1-2%)(*39*). One possible explanation for this apparent discrepancy may come from the fact that WSD habits, often in combination with chronic over-nutrition and sedentary lifestyles, have been associated with a state of chronic metabolic inflammation, termed ‘metaflammation’(*40*). In particular, a link between consumption of a diet high in fat and sugar and Paneth cell dysfunction was recently demonstrated(*41*). Moreover, in a recent report by J.C. ad L.H.A., a purified mouse diet that mimics WSD habits and underlies the increased risk of colon cancer (NWD1)(*42*) was shown to induce a low degree of chronic intestinal inflammation and other mechanisms that define pathogenesis of human IBD(*43*). Therefore, we hypothesized that etiologic drivers of colon cancer other than IBD, including widespread western-style dietary habits, may underlie dedifferentiation and tumor onset mechanisms similar to those observed upon inflammation.

To provide support for this hypothesis, we first fed C57BL6/J mice for 3 months with the western-style (NWD1) and control (AIN76A) diets and compared the transcriptional response of Paneth cells with that obtained upon DSS administration (Fig. 4a). We examined genes upregulated upon inflammatory stimuli (DSS signature; see Methods), which showed variable but overall increased levels in NWD1-fed mice when compared to those on the control diet (Fig. 4b). Indeed, GSEA confirmed that the DSS signature was associated significantly with Paneth cells exposed to NWD1 (NES 2.99, Padj < 0.001), indicating that western-style dietary habits trigger an inflammatory-like response in Paneth cells (Suppl. Fig. 9a). At the gene ontology level, western-style diet activated signaling pathways related to cell cycle (G2M Checkpoint) and proliferation (mitotic spindle, Myc targets; Suppl. Fig. 9b), suggesting that Paneth cells re-enter the cell cycle upon long-term exposure to western-style diet, similar to what is observed upon DSS-driven inflammation(*11*).

**Figure 4.**
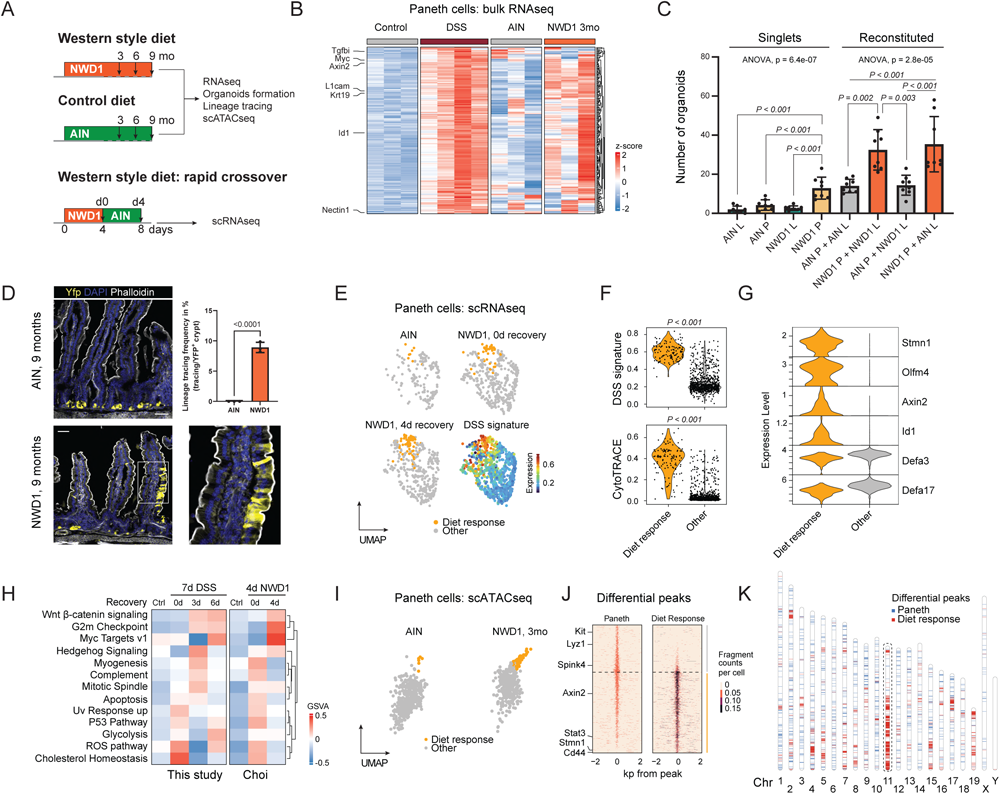
Western-style diet triggers an inflammatory response leading to Paneth cell dedifferentiation. **a.** Schematics of the experimental approach designed to investigate the consequences of short-and long-term exposure to western-style diet (NWD1) vs. control (AIN76A) diets. **b**. Heatmap showing z-scored DSS signature (DSS vs. Control; Padj < 0.05, Log2FC > 0.25) in Paneth cells exposed to DSS or NWD1. **c**. Organoid multiplicities derived either from single ISCs or PCs, and from reconstituted doublets (L: Lgr5+ ISCs, P: Paneth cells). Pooled data from N = 4 independent experiments. P values were calculated using one-way ANOVA and Tukey tests for group comparisons. Error bars depict standard deviations from the mean. **d**. Representative image of lineage tracings from a NWD1-fed Lyz1-Yfp mouse. Scale bar: 50 μm. P value depicts result of the student t-test and error bar represents the standard deviation. Data from N = 3 mice. **e**. UMAP showing Paneth cells from AIN76A-and NWD1-fed mice (N = 3 mice per condition). The DSS signature portrayed on UMAP embedding, highlights a subcluster of Paneth cells responsive to the NWD1 diet. **f**. Violin plots showing different levels of the DSS signature (top) and CytoTRACE score (bottom) between the PCs responsive to the NWD1 diet and other Paneth cells. P values depict significance values of Wilcoxon test. **g**. Violin plots representing marker genes of PCs responsive to the NWD1 diet, showing co-expression of stem and secretory markers. **h**. Heatmap visualization of gene set variation analysis, indicating pathways that are activated in Paneth cells after exposure to DSS or NWD1. **i**. UMAP plot of Paneth cell subset in scATAC data set of AIN (N = 2) and NWD1 (N = 2) treated mice. **j**. Heat map listing differential peaks between Diet response Paneth cells and other PCs. **k**. Ideogram displaying the distribution along the mouse chromosomes of the differential peaks observed upon diet response (red) cell when compared to those characteristic of Paneth cells (blue).

In view of the previously reported acquisition of stem-like features by PCs upon inflammation(*11*), we next employed the organoid reconstitution assay (ORA)(*10, 44*) to assess whether similar effects are exerted by the western-style diet. We first co-incubated Paneth and *Lgr5*^+^ cells from the AIN76A-and NWD1-fed mice in all 4 combinations. As shown in Figure 4c, Paneth cells from NWD1-fed animals significantly improved organoid formation independently of their reconstitution with *Lgr5*^+^ cells from NWD1-or AIN76A-fed mice, possibly indicative of a paracrine effect enhancing the well-established niche (ISC-supporting) role of Paneth cells. However, single (i.e. non reconstituted with *Lgr5*^+^ CBCs) Paneth cells from NWD1-fed mice formed organoids more efficiently when compared with either PCs from AIN76A-fed mice or with *Lgr5*^+^ ISCs from both groups of animals. These *ex vivo* results were further validated by lineage tracing analysis of Paneth cells in R26^LSL-YFP^*Lyz1*^CreERT2^ NWD1-fed mice that revealed extended YFP-labeled ribbons thus confirming their de-differentiation and acquisition of stem-like features induced by the western-style dietary cues (Fig. 4d).

To zoom in on the primary transcriptional response of Paneth cells to NWD1, we took advantage of the scRNAseq data generated by J.C., L.H.C. and collaborators immediately upon exposure to NWD1(*43*)(Fig. 4a). Within 4 days of switching mice to the NWD1 diet, a subset of Paneth cells became apparent whose transcriptional profile strongly associated with the DSS signature (labeled as “diet response cells” in Fig. 4e). Mirroring our previous observations obtained immediately upon DSS inflammatory stimulus, these WSD-responsive cells increased their transcriptomic diversity as measured by CytoTRACE(*45*) (Fig. 4f; see Methods). Already after the short exposure to NWD1, the diet-responsive cells acquired stem cell markers while retaining some secretory features (Fig. 4g). Comparative pathway analysis between the transcriptional response of Paneth cells to DSS and NWD1 revealed similar upregulation of Wnt, Myc, Hedgehog, and G2M checkpoint signaling pathways (Fig. 4h).

As the observed WSD-driven gene expression changes are likely exerted through epigenetic modifications, we next analyzed scATAC data obtained in the framework of the same Choi et al. (2023) study(*43*). Similar to what observed by scRNAseq analysis, we identified a group of NWD1 diet-responsive Paneth cells with significant epigenetic modifications including a main cluster (52%) on mouse chromosome 11 (synthenic with human chromosomes 17 and 5) that encompasses a considerable fraction of genes encoding for members of the Wnt, Pi3k/Akt and cell cycle pathways (Fig. 4i-k, Suppl. Fig. 9c-f). The latter were previously shown by our laboratory to underlie Paneth cell dedifferentiation and acquisition of stem-like features upon DSS-driven inflammation(*11*).

In order to assess the effects of the NWD1 diet on the mouse colon, we looked for proliferating Paneth-like cells, also known as deep crypt secretory (DCS) cells(*7, 10*). Inflammation-driven cell cycle activation in these allegedly post-mitotic lineages was previously observed in small intestinal Paneth cells of DSS-treated mice and of Crohn’s disease patients(*11*). By co-staining colonic tissues with the secretory lineage marker WGA (wheat germ agglutinin)(*46*) and Ki67, a significant increase in the number of proliferating secretory cells located at the crypt bottom was observed both in mice fed with the NWD1 diet and, as a positive control, in those administered DSS in the drinking water (Fig. 5a-c).

**Figure 5.**
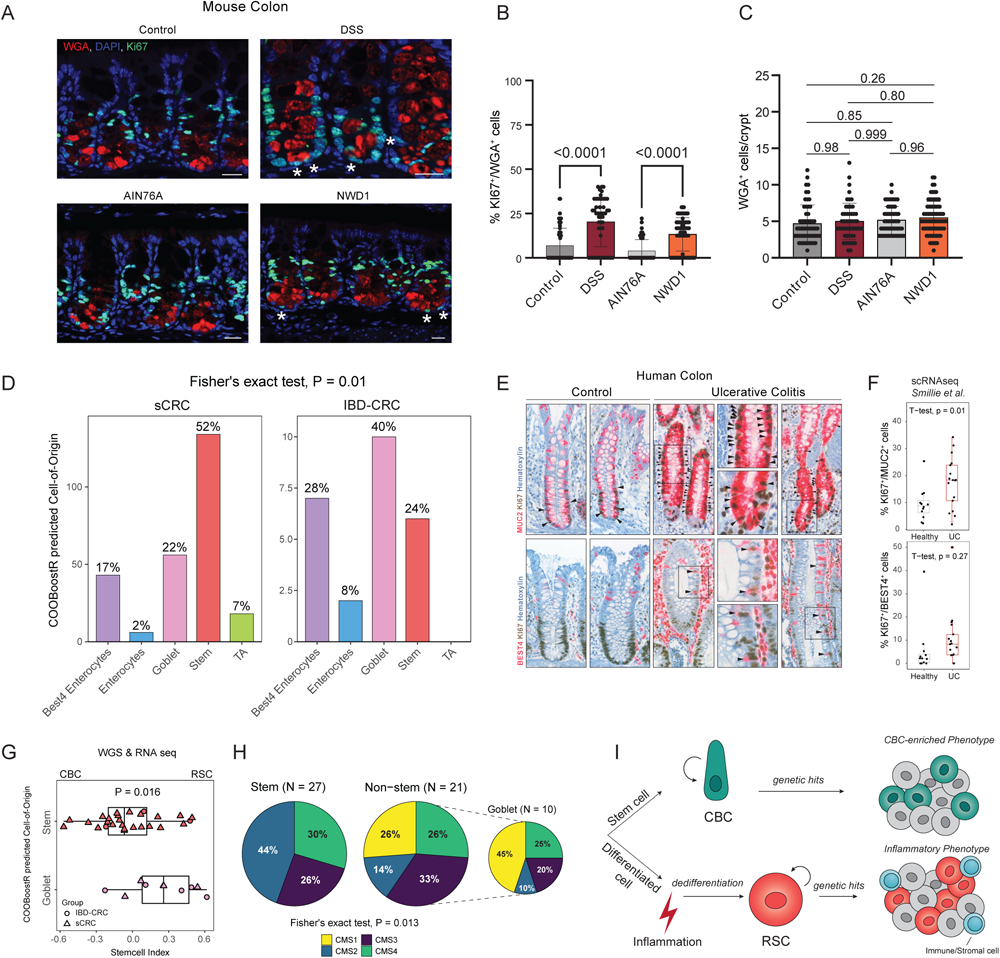
Inflammation activates distinct cells of origin in the colon. **a.** Colonic tissues from either untreated mice or from mice administered 3% DSS for 7 days, as well as from mice fed with the AIN76A or NWD1 synthetic diets for 3 months were analyzed for the presence of proliferative deep crypt secretory (DSC) cells. WGA (wheat germ agglutinin) was used to stain DSC cells(*46*), and Ki67 to mark proliferative cells. Tissues were counterstained by DAPI (nucleus). Asterisks mark WGA/Ki67 double positive cells. **b-c**. Quantification of WGA^+^/Ki67^+^cells (**b**) and total WGA^+^ cells (**c**) in the lower colonic crypt of the mice, as shown in (**a**). Scale bar = 20 μm. A minimum of 50 crypts from 3 different mice were analysed. **d**. Barplot showing the predicted cell-of-origin in a cohort of IBD-CRC (N = 25) and sCRC (N = 257)(*35*) based on the COOBoostR computational approach(*49*) (see Methods). P value depicts result of the Fisher exact test. **e**. MUC2, BEST4, and Ki67 IHC analysis of colonic tissues obtained from controls and IBD patients. Asteriks mark examples of double positive cells. **f**. Boxplot showing percentage of cycling (Ki67^+^) MUC2^+^ and REG4^+^ cells in Ulcerative Colitis patients and controls. scRNAseq data are from Smillie et al.(*51*). Positivity was defined per cell by the presence of at least one read for that particular marker. Subsequently, cells were aggregated per patient to calculate percentages. **g**. Box plot denoting differences in the stem cell index based on stratification of predicted cell-of-origin in a subset of IBD-CRC and sCRC cases for which RNAseq data were available(*35*). P value depict the result of the t-test. **h**. Mapping of Consensus Molecular Subtypes (CMS) on tumor samples stratified according to their predicted cell-of-origin. P value depicts result of the Fisher exact test. **i**. Graphic abstract of the model arising from the present study. Colon cancer can be initiated either from stem (CBC) or differentiated cells, the latter in response to inflammatory cues. RSC reprogramming is activated in support of the regenerative response. During this process, actively dividing RSCs expand the cell targets for tumor initiation and progression, leading to an alternative route to tumorigenesis earmarked by an inflammatory phenotype.

### Secretory lineages as the cell of origin of human colon cancer

Collectively, our results reveal an alternative bottom-up route to intestinal tumorigenesis originating from Paneth cells in the murine small intestine and, allegedly, from secretory lineages in the human colon triggered by inflammatory cues as in IBD or through western-style dietary factors. To validate the relevance of our mouse study in patient-derived colon cancer, we took advantage of novel computational methods(*47–49*) developed to predict the cell of origin of tumors by matching the mutational density along the cancer genome with the profiles of epigenetic modifications characteristic of normal cell types (Suppl. Fig. 10a)(*50*). While tumors from patients with an IBD history have a similar genome-wide tumor mutational burden compared to their sporadic counterparts (Suppl. Fig. 10b), the presence of regions (“genomic windows”, see Methods) that are differentially mutated between sCRC and IBD-CRC is suggestive of alternative mutational patterns (Suppl. Fig. 10c-d). Hence, we compared by COOBoostR(*49*) (see Methods) the individual mutational landscapes with the epithelial cell types of the colon to compute the putative cells of origin. As shown in Fig. 6a, while the majority of the sporadic colon cancers appear to originate from stem cells (52%), among the IBD-related cases goblet (40%) and Best4 (28%) cells represent the prevalent cells of origin. Of note, a substantial proportion of the sporadic cases is also predicted to originate from goblet (22%) and Best4 cells (17%). Best4 cells, named after the specific expression of the bestrophin 4 gene (*BEST4*), form a newly identified and yet only partially characterized intestinal epithelial lineage with dual absorptive and secretory features(*51–53*). IHC and scRNAseq analysis in a cohort of ulcerative colitis patients confirmed the presence of actively proliferating goblet and Best4 cells (MUC2^+^ and BEST4^+^; Fig. 5e-f), indicative of the primary effects of inflammation in these otherwise post-mitotic cells, preceding their acquisition of stem-like features.

These results are indicative of the fraction of sporadic colon cancers whose expression profiles are reminiscent of the PC-derived tumors and IBD-CRCs (24.7%; Figure 3j). Furthermore, the availability of RNAseq data from a subset of the patient-derived tumors allowed us to confirm the prevalence of RSC-like expression profiles and the high proportion of CMS1 and 4 cases among the sporadic and IBD cases predicted to originate from non-stem, and in particular goblet, cells (Fig. 5g-h, Suppl. Fig. 10e).

*Lgr5*^+^ stem cells have been established as the origin of intestinal tumors(*1*). However, whether the same holds true in the context of inflammation has been challenged by their loss of lineage tracing capacity and multipotency upon a broad spectrum of tissue injuries ranging from DSS-driven inflammation(*11*) to γ-irradiation(*9, 22*), and western-style diet(*54*). Our results show that Paneth cell de-differentiation and tumor onset in the mouse small intestine models sporadic colon cancer not only in IBD patients but also in a substantial proportion of the cases predicted to originate in differentiated cell types and not in resident ISCs. Both the differential gene expression profiles and inflammatory and stromal features of the resulting tumors warrant a novel stratification of colon cancer for improved clinical management. Preliminary investigations (*data not shown*) revealed that candidate markers differentially expressed between PC-and *Lgr5*^+^-initiated mouse tumors such as DCLK1 and HOXB9 were not equally discriminative among patient-derived colon cancers. More systematic and high-throughput screening approaches will likely lead to a novel stratification of the cases based on their cell of origin and specific etiologic factors such as western-style diets and chronic inflammation.

Overall, the increased colon cancer risk exerted by inflammation may reflect on a ‘trade-off’ effect where the chronic nature of the injury continuously stimulates de-differentiation of committed lineages into stem-like cells in support of the tissue’s regenerative response, thus resulting in the enlargement of the pool of potential targets for tumor onset. Likewise, the same inflammation-inducing risk factors are bound to affect tumor progression towards malignancy thus resulting into a distinct group of colon cancers with specific prognosis and response to therapy (Fig. 5i).

Relative to the debate on the contribution of extrinsic risk factors versus the rate of stem cell division to cancer development(*55, 56*), our results indicate that colon cancer etiologic factors such as inflammation and poor dietary habits are likely to result in quantitative and qualitative alterations of the stem cell niche which ultimately predispose to neoplastic transformation. The resulting subset of tumors follow distinct evolutionary paths when compared to Wnt/Myc-driven *Lgr5*-derived tumorigenesis, and are characterized by an inflammatory tumor phenotype more prone to infiltration from the tumor micro-environment.

## Methods

### Mice

The following inducible Cre strains encompassing were employed: *Lgr5*^CreERT2-EGFP^ (#008875, Jackson Lab(*15*)) and p*Lys*^CreERT2^ (kind gift from the Clevers Lab(*16*)). For lineage tracing experiments, mice were crossed with R26^LSL-YFP^ mice (#006148, Jackson Laboratories) or R26^LSL-tdTomato^ mice (#007908, Jackson Laboratories). To target specific mutations in PCs, the above strains were further crossed with *Apc*^15lox^ (#029275, Jackson Laboratories)(*12*), *Kras*^LSL-G12D^ (#008179, Jackson Laboratories)(*13*), and *Trp53*^flox^ (#008462, Jackson Laboratories)(*14*). For ablation experiments Lgr5^DTR-EGFP^ mice were bred with c-Kit^CreERT2^ and R26^LSL-tdTomato^ animals. C-recombinase was activated by intraperitoneal injections of Tamoxifen (1 mg dissolved in 100% EtOH and subsequently in sunflower oil; #T5648 and #S5007, Sigma) once or three times in 4 days. Three days after Tamoxifen injection mice were treated once with diphteria toxin (DT) in order to ablate Lgr5 positive cells. To induce acute intestinal inflammation, mice were administered 2-3% dextran sodium sulfate (DSS) in their drinking water for 7 days (#0216011050, MP Biomedicals). DSS-driven inflammation and the corresponding mechanisms underlying the consequent PC dedifferentiation were described in previous studies(*11, 57, 58*).For diet experiments, mice were fed with AIN76A or NWD1 diet(*54*) and collected at various time points (4/8 days, 3/6/9 months). The formulation and composition of the AIN76A and NWD1 diets are as outlined in Newmark et al. (Research Diets, New Brunswick, NJ)(*42*). In brief, the western diets are based on AIN76A control diet (American Institute of Nutrition76A). NWD1 was adjusted to provide higher fat, and lower vitamin D3, calcium donors to the single carbon pool and fibre that produces mouse consumption levels similar to those common in segments of western societies with high incidence of colorectal cancer. All mice here employed were inbred C57BL6/J.

For all experiments, mice were randomly assigned to experimental groups after matching for gender, age of 8-12 weeks, and genotype. All protocols involving animals were approved by the Dutch Animal Experimental Committee and in accordance with the Code of Practice for Animal Experiments in Cancer Research established by the Netherlands Inspectorate for Health Protections, Commodities and Veterinary Public Health. Animals were bred and maintained in the Erasmus MC animal facility (EDC) under conventional specific pathogen-free conditions.

### Lineage tracing

*Lys*^CreERT2/R26LSL-YFP^ mice were injected three times with Tamoxifen (1 mg, i.p.) on consecutive days. One week after the last Tamoxifen injection, intestinal tissues were harvested for lineage tracing analysis. To this aim, tissue samples were first dissected and washed with PBS, and then fixed for 2 hours at 25°C with 4% buffered formaldehyde solution (Klinipath). Tissues were cryo-protected in 30% sucrose (Sigma) overnight at 4°C, embedded in OCT (KP cryocompound, Klinipath), frozen on dry ice, and sectioned at −20°C. Tissues were cut in 4-8 μm thick sections. These sections were incubated in PBS containing Alexa 568 Phalloidin (1:100; Invitrogen) or Alexa 633 Phalloidin (1:100; Invitrogen), and DAPI (Sigma) for 30 minutes at 25°C and washed in PBS-T. Tissues were mounted in Vectashield Mounting Medium (Vector Labs) and imaged with a LSM700 confocal microscope (Zeiss). Images were processed with ImageJ. Lineage tracing frequency was quantified by dividing the number of intestinal crypt/villus axes containing Yfp^+^ ribbons (encompassing at least 5 Yfp^+^ labelled cells) by the total number of counted Yfp labelled crypts. Lineage tracing analysis was performed on 3 mice for each diet type; in each mice at least 60 YFP labelled crypts were analyzed.

### Immunohistochemistry on mouse tissues

Tissues were fixed in 4% PFA overnight at 4°C and embedded in paraffin. The 4 µm sections were dewaxed with Xylene and hydrated in consecutive rounds of 70% and 100% EtOH. Antigen retrieval was performed in the 2100 Retriever pressure cooker (BioVendor) at pH 9 with Tris-EDTA buffer. After a 10 minutes incubation at RT with 3% hydrogen peroxidase, tissues were blocked with 5% skim milk powder (Milipore) in PBS-Tween. The following primary antibodies were employed by overnight incubation at 4°C: β-catenin, #610154 (BD Biosciences); γ-H2AX, #9718 (Cell Signaling); Gfp, #A-11122 (ThermoFisher); Olfm4, #D6Y5A (Cell Signaling); Lyz1, #A0099 (Dako); Dclk1, #ab37994 (Abcam); Muc2, #sc-15334 (Santa Cruz); Chga, #NB120-15160 (Novus Bio); Ecad, #610182 (BD Biosciences); Cd3, #ab5690 (Abcam); F4/80, #D2S9R (Cell Signaling); αSMA, #ab5690 (Abcam). Slides were washed twice with PBS-Tween and incubated for 30 minutes at room temperature with the Rabbit/Mouse EnVision kits (K4001/K4007, Dako). Slides were counterstained with hematoxylin and, dehydrated with subsequent 70% and 100% EtOH and mounted with Pertex (00811, Histolab). Whole slides were scanned with the Nanozoomer (Hamamatsu) and analyzed with NDP viewer v2 (Hamamatsu).

## Immunohistochemistry analysis of human tissues

Chromogenic double labelling was performed on 4-µm thick whole slide sections from FFPE tissue blocks, on a validated and accredited automated slide stainer (Benchmark ULTRA System, VENTANA Medical Systems, Tucson, AZ, USA) according to the manufacturer’s instructions. Briefly, following deparaffinization and heat-induced antigen retrieval for 40 minutes at 97°C, the tissue samples were incubated with KI67 (MIB-1 antibody, #790-4286; Ventana) for 32 min at 37°C, followed by Ultraview detection (#760-500, Ventana). After the stripping step, MUC2 (CCP58, #M7313, Dako) or BEST4 (HPA058564, Sigma-Aldrich) was incubated for 32 min at 37°C and followed by detection with Ultraview Red (#760-501, Ventana) Counterstain was done by hematoxylin II for 12 min and a blue colouring reagent for 8 min. Each tissue slide contained a fragment of FFPE pancreas or tonsil as an on-slide positive control.

### Immunofluorescence

For cryo-sections, tissue samples were first dissected and washed with PBS, to then be fixed for 2 hours at 25°C with 4% buffered formaldehyde solution (Klinipath). Tissues were cryo-protected in 30% sucrose (Sigma) overnight at 4°C, embedded in OCT (KP cryocompound, Klinipath), frozen on dry ice, and sectioned at −20°C. Tissues were cut in 4-8 μm thick sections. Immunofluorescence was performed using antibodies against Olfm4 (1:200; CST), Mki67 (1:200; Novus biologicals), Lyz1 EC3.2.1.17 (1:1000; Dako), and c-Kit (D13A2)(1:100; CST). Tissue sections were washed and blocked with 5% BSA in PBS-Tween for 1 hour to then be incubated with the primary antibody overnight at 4°C. Slides were washed twice with PBS-Tween and slides were incubated for 2 hours with Goat anti-Rabbit IgG (H+L) secondary Antibody, Alexa Fluor® 488 conjugate, (1:250; Life Technologies) and Wheat Germ Agglutinin (WGA)^TM^ 647 conjugate (1:500, ThermoFisher Scientific). Tissues were counterstained with DAPI (Sigma) and where indicated with F-actin (#SC001, Spirochrome) for 30 min. at 25°C and washed in PBS-T. Tissues were mounted in VECTASHIELD HardSet Antifade Mounting Medium (Vector Labs) and imaged with a Zeiss LSM700 confocal microscope.

### In situ hybridization

In situ hybridization was performed by RNAscope^TM^ Multiplex Fluorescent V2 Assay (ACDBio) technology in combination with the RNA-Protein Co-Detection kit (ACDBio), according to the manufacturer’s instructions. In brief, 4 µm paraffin embedded sections were dehydrated, blocked for 10 minutes at room temperature with H2O2, and processed with 1X Antigen Retrieval co-detection buffer for 15 minutes at 99°C. Sections were incubated with Ecad (1:250, #610182, BD Biosciences) overnight at 4°C. Hybridization was performed with a probe targeting mouse *Clu* (Cat#427891, ADCBio), and the probe signal was developed with Vivid dye 520 nm (1:1500). Sections were counterstained by DAPI and anti-mouse Alexa 633 (1:250, Invitrogen) and mounted with ProLong^TM^ Gold Mountant (ThermoFisher). Samples were imaged on the Stellaris 5 Confocal Microscope (Leica) and images were processed with ImageJ.

### Image analysis

Images were scanned by Nanozoomer and imported in QuPath(*59*) as IHC images. Within intestinal tumors, regions of interest (ROI) were randomly assigned and consisted of 300-600 cells. Next, the ROIs were processed with positive cell selection plugin using default settings based on Cell DAB OD mean. The percentage of positive cells per ROI was exported and visualized in R, where statistical tests were performed. Fluorescent images were processed with ImageJ.

### Quantification of Ki67/WGA cells in murine colon

The percentage of Ki67^+^/WGA^+^ double positive cells was quantified by dividing double positive cells by the total amount of WGA^+^ in the lower half of the colonic crypt. Only clearly detectable double positive cells, as determined by the analysis of z-stacks were counted. For each quantification, at least 30 crypts of at least 3 experimental mice were employed.

### Quantitative real-time PCR

RNA was converted into cDNA using the High Capacity RNA-to-cDNA kit (Applied Biosystems). Samples were run in duplicates on the 7500 Real Time system (Applied Biosystems). Quantitative PCR was performed using the TaqMan assay (Applied Biosystems) according to instructions of the manufacturer. Samples were normalized to the beta-actin (*Actb*) house-keeping gene. The gene-specific Taqman probes were: Mm02619580_g1 (*Actb*), Mm00443610_m1 (*Axin2*), Mm00657323 (*Lyz1*), Mm01545303 (*Dclk1*). Data were processed and visualized in R and statistical tests were employed to assess significance with one-way ANOVA and Tukey post hoc test.

### Isolation of mouse intestinal crypts and cell dissociation

Mouse small intestines were flushed with cold PBS, removed from fat and scraped with a glass slide to remove intestinal villi. Resected sample were then sectioned into 5-10 mm segments and washed with cold PBS before incubation for 30 minutes at 4°C in cold PBS supplemented with 6 mM EDTA. After an additional PBS wash, crypts were detached from the muscle layer by 4 rounds of harsh pipetting with cold PBS. Crypts were treated for 10 minutes at room temperature with DNAse (2000 U/ml-1, Thermo Scientific) in Advanced DMEM/F-12 medium and passed through a 70 µm strainer. Purified crypts were dissociated with 1 mL of pre-warmed TryplE and DNAse for 3 minutes at 37°C. Single cells were pelleted in 10 mL ADF, resuspended in 3 mL 5% FCS HBSS and manually counted.

### Fluorescence activated cell sorting

Cell suspensions were blocked with TruStain Fc blocking reagent (Biolegend) for 10 minutes at 4°C. After washing with 5% FCS HBSS, cells were stained for 30 minutes at 4°C in 100 µl with Lin antibodies (CD31-BV421 #563356, CD45-BV421 #563890, TER119-BV421 #563998, BD Biosciences), CD24-APC (#1109070, Sony Biotechnology), and cKit-PE (#105808, Biolegend). For scRNAseq experiments, the antibody mix was supplemented with 2 µl Hashing Antibody (Biolegend TotalSeq^TM^ Hashtag antibodies: #155831, #155833, #155837, #155843, #155849) to label the cells with oligo barcodes. After 3 washes with 3.5 mL 5% FCS HBSS, cells were filtered with a 40 µm strainer and resuspended in 1 mL 5% FCS HBSS with 1 µg/mL DAPI (D9542, Sigma-Aldrich). FACS analysis and sorting were performed using a FACSAriaIII (BD Biosciences). DAPI and BV421-conjugated antibodies were detected using a 405nm laser and 450/40 BP filter. GFP and BB515-conjugated antibodies were detected using a 488nm laser and 502LP + 530/30 BP filters. tdTomato and PE-conjugated antibodies were detected using a 561nm laser and 582/15 BP filter. APC-conjugated antibodies were detected using a 633nm laser and 660/20 BP filter. Samples were sorted for epithelial (Lin^neg^), Paneth-enriched (SSC^hi^CD24^hi^cKit^hi^), and traced cells (Yfp^hi^/td-Tomato^hi^) according to previously established gates(*9–11*). To optimize the sorting for single cell experiments, the initial single cell gate was followed by a more stringent, population specific single cell gate(*9*). FACS strategy was visualized with FlowJo V10.

### Organoid Reconstitution Assay

Whole crypts were extracted from the small intestine of *Lgr5*-EGFP mice fed for 3 month (from weaning) with either AIN76A or NWD1 diets(*54*). After cellular dissociation and FACS, single *Lgr5*^+^ and Paneth cells were sorted in separate Eppendorf LoBind Tubes. Reconstitution of *Lgr5*-EGFP^hi^ stem cells with PCs (SSC^hi^CD24^hi^cKit^hi^) was performed by co-pelleting sorted cells at 300 g for 5 minutes in Eppendorf LoBind Tubes, and by incubating them for 15 minutes at 25°C as previously described(*10, 44*). The cells were then resuspended in 30 μL Matrigel (Corning), and cultured in 96 Well Flat Bottom dishes (Corning). Additionally, Paneth and Lgr5 positive cells were plated singularly where indicated and cultured in standard ENR medium(*60*) supplemented with 10 μM Y-27632 (Sigma) and 1 mM Jagged-1 (AnaSpec). Cells were plated in triplicates and the generated organoids where counted 9 days after plating. Experiments were repeated at 3 independent moments.

### Droplet-based scRNAseq

Sorted single cells were mixed in equal sample proportions to a pool of 25-50k cells in 5% FCS HBSS to a final concentration of ±1000 cells/µl. The cell pool was recounted manually with Trypan blue and between 10-15k cells were loaded into a 10X Genomics Chip (Next GEM Chip G). The Gel Bead-In EMulsions were generated with the Chromium Controller (10X Genomics) using Single Cell 3’ v3 chemistry with FeatureBarcoding. After generation of cDNA and quality check with Bioanalyzer (Agilent), libraries were sequenced with the Novaseq 6000 (Illumina) till a depth of 20-40k reads/cell. Raw data was processed with CellRanger count (cellranger-7.0.0)(*61*) and aligned to the mm10 mouse reference genome with the addition of the transgenes Yfp and Td-tomato.

### Analysis of scRNAseq: inflammation and genetic targeting of Paneth cells

For each single cell run, filtered gene-cell matrices were imported in R using Seurat (v 4.1.1)(*62*). After a centered log ratio transformation (CLR) of the hashtag oligo counts, cells were assigned back to their mouse of origin using HTODemux (positive quantile = 0.99). Cells assigned as doublets or negative for any of the barcodes were filtered from the data. Cells from mice with different gender but same genotype and experimental group were further separated based on the expression *Xist*. After preprocessing, the five runs were merged into a single Seurat object. To remove putative doublets from the sorting, doublets were simulated and called by doubletfinder_v3 (nExp = 0.01)(*63*). After removal of doublets, low quality cells were removed by filters based on the percentage of mitochondrial genes (> 10%), on nFeature_RNA (< 200), and nCount_RNA (< 500). Next, data were integrated with the single cell transform (SCT) pipeline using reciprocal PCA dimension reduction to find integration anchors. Integrated data was further processed by dimension reduction with Uniform Manifold Approximation and Projection (UMAP) on the first 30 principal components. Cells were clustered using shared nearest neighbor (SNN) modularity optimization (resolution = 0.6) and annotated according to canonical marker genes. Traced cells were defined as cell with a non-zero value for either td-Tomato or Yfp. Signatures were evaluated using the Addmodulescore function and visualized in experimental groups or cell clusters using Violin Plots. Pathway level evaluation was done using the GSVA package(*64*) on summarized expression profiles using Average Expression by cell cluster.

### Bulk RNAseq of intestinal tumors

Intestinal tumors were macro-dissected from intestinal tissues and cut into 1-2 mm small fragments which were then re-suspended in 500 µl TRIzol (#15596026, Ambion) for RNA isolation. RNA quality was first evaluated by Nanodrop ND-1000 (ThermoFisher) and further purified with the TURBO DNA-free Kit (Invitrogen). Samples were sequenced with a read length of 150 bp (PE150) using the DNA nanoball (DNB-seq) protocol till a depth of 25M reads/sample (BGI, Hong Kong). Adapter trimming and sequencing QC was performed with SOAPnuke software(*65*).

### Analysis of Lyz1/Lgr5-derived mouse tumors: bulk RNAseq

The FASTQ files were aligned with STAR (v2.7.9a) to the mm10 mouse reference genome with Ensemble gene annotations(*66*). After alignment, Sambamba (0.8.0) was applied to mark duplicates and perform flagstat quality checks(*67*). Next, Subread (2.0.3) was used to count the primary strand-specific alignment(*68*). Downstream analysis was performed in R using the DeSeq2 package (v 1.34.0)(*69*). In brief, counts were log2 normalized and used for principal component analysis on the top 500 variable expressed genes. Differential expression analysis was done following the DeSeq pipeline with Negative Binomial GLM fitting and Wald statistics and genes were filtered using log_2_FC >1.5, P_adj_ < 0.01. Gene set enrichment analysis was done with the fgsea package (v 1.20.0) using the HallMarks gene modules from the molecular signature database(*36*). Gene set variation analysis was performed using the GSVA package (v 1.42.0)(*64*). The relative proportion of RSC/CBC stem cell types was computed with the ISC index using default settings(*29*).

### Analysis of Paneth cells upon exposure to inflammation and western-style diet: bulk RNAseq

The FASTQ files were removed from TrueSeq adapters using Trimmomatic (v0.36). STAR(*66*) was used to align the reads to the mm10 reference genome using Gencode annotation release M15 (GRCm38.p5). Mapping quality was assessed with Sambamba Flagstat statistics. Count files were generated using FeatureCounts (subread) and further processed in R using the DeSeq2 package. Differential expression analysis were performed between the groups ‘NWD1 diet’ (n = 3) vs. ‘AIN76 diet’ (n = 3) with the Wald-test. A heat map was produced using z-score scaled data and visualized with the ComplexHeatmap package (v2.12.1).

### Analysis of IBD-CRC and sCRC RNAseq cohort

RNA-seq material was previously described in Rajamäki et al.(*35*) In brief, 64 CRCs entered HiSeq LncRNA-Seq library preparation and paired-end sequencing using Illumina HiSeqXTen. Salmon (version 0.12.0; quant mode with ValidateMappings) was used to map raw sequences onto the human transcriptome (Ensembl release 79). Gene-level quantification was done with DESeq2 (v1.18.1) followed by a variance stabilizing transformation and Limma (v3.34.9) correction of sequencing batch effects(*69*). Only microsatellite stable (MSS) tumors were included in the analyses. Differential gene expression was calculated as ANOVA between 38 MSS sCRCs and 14 MSS IBD-CRCs. GSVA (v1.42.0)(*64*) analyses were ran using default options (Gaussian kernel) and predefined gene sets from the molecular signature database(*36*). Conensus Molecular Subtypes were mapped to the samples with the random forest classifier of CMSClassifier R package.

### Cell of origin predictions from IBD-CRCs/sCRCs

#### Processing of Whole-Genome Sequencing (WGS) data of CRCs

DNA was isolated from fresh-frozen tumor, normal colon, or blood. Paired tumor-normal WGS data was generated from altogether 25 MSS IBD-CRCs and 257 MSS sCRCs. Libraries were prepared using TruSeq Nano DNA HT or TruSeq PCR-Free Kit (Illumina), followed by paired-end sequencing using Illumina platform (HiSeqXTen/HiSeq2000). More details are given in Rajamäki et al.(*35*). Sequence alignment to the GRCh38 reference genome, other data preprocessing steps, and somatic variant calling were performed with GenomeAnalysisToolkit GATK4 best practices workflow (version 4.0.4.0)(*70*). Mutations were filtered based on allele fraction (AF) to extract clonal somatic variants. A genome-wide mean and standard deviation was calculated for each tumor; somatic variants were then filtered to a minimum AF of one standard deviation below the mean. On average, 85% of variants per tumor (IQR 83%-87%) were kept-in as clonal somatic variants for the subsequent analysis. Similar to previous work(*47, 48*), the human reference genome (hg38) was processed in non-overlapping windows of 1 million base pairs. We excluded window-regions with <92% of bases within uniquely mapped 36-mers, regions overlapping telomeres and centromeres and regions outside autosomes. Altogether 2387 windows remained for the analysis. To identify differentially mutated windows, patients were normalized by the average of their windows and IBD-CRCs vs sCRCs were compared with an Wilcoxon Rank Sum test (P < 0.01).

#### Processing of scATAC data from human colon

Normal colon (N = 9) samples from Becker et al.(*50*) were obtained from the public domain under GEO identifier GSE201349. Signac(*71*) was used for processing and the FeatureMatrix function was used to generate a count matrix based on the previously described 2387 windows. Cell labels were retrieved from the original publication and reduced to 6 major classes by the aggregation of clusters from the same cell type. To generate the embedding, batch correction was performed with IntegrateData, and UMAP was computed on the 2:30 latent semantic indexing (LSI) components. To generate the chromatin mark features per cell type, the counts from the scATAC data set were grouped with the AggregateExpression function and equalized to 10M reads per cell type.

#### Predictions with COOBoostR

COOBoostR(*49*), an extreme gradient boosting-based (XGBoost) machine learning algorithm designed for the prediction of cells of origin, was used to predict the putative cells of origin for each individual sample from the WGS cohort (N = 282 tumors) based on the epigenetic profiles from the human colonic scATAC data. We ran the algorithm with default values from XGBoost for the learning rate (mEta = 0.3) and the maximum depth of the trees (mdepth = 6). Tumor predictions were aggregated on cohort level, and IBD-CRC/sCRC were compared with a Fisher’s exact test. For a subset of patients (N = 48 tumors), WGS and RNAseq was available. Hence, for these tumors, CMS labels and signatures from the gene set variation analysis (GSVA) were compared based on stratification by putative cell of origin.

### Analysis of human CRC bulk RNAseq cohort

Log2 normalized expression of the meta cohort described in Guinney et al.(*38*) was obtained from the R2 Genomics Analysis and Visualization Platform and subsequently processed in R. Gene set variation analysis was performed with the GSVA package using the Gaussian kernel(*64*). Subsequently, correlations of the gene signatures (Suppl. Table 4) were visualized with ggcorr using Pearson Correlation. Next, samples were classified as colitis-like, sporadic-like or as intermediate based on their GSVA scores for the IBD-CRC and sCRC signatures. Using this classification, samples were visualized with their annotated Consensus Molecular Subtype (CMS). Survival analysis was performed with the survival and survminer packages in R. Multivariate analysis was performed with the log-rank test and pairwise comparisons were fitted with a Cox proportional hazards regression model.

### Analysis of human scRNAseq cohorts

#### Colorectal cancer data set

Filtered count matrices were retrieved from the gene expression omnibus (KUL cohort: GSE144735; SMC cohort: GSE132465)(*37*) and processed in R with Seurat. After normalization and scaling of the Seurat object, gene expression was summarized per cell type using AverageExpression based on the overlapping differentially expressed genes of murine (Paneth vs. *Lgr5*) and human (IBD-CRC vs. sCRC) tumor groups. The average expression was z-scored and visualized as a heat map using the ComplexHeatmap package.

#### Ulcerative Colitis data set

Data from Smillie et al.(*51*) was processed with the Seurat(*62*) workflow. A subset was taken encompassing the epithelial cells according to annotation from the authors. Cells were defined as positive when they had non-zero counts for that particular gene. Cells were aggregated on patient level and a t-test was performed to compare number of double and single positive cells across the UC (N = 17) and control group (N = 12).

### Analysis of mouse intestinal single cell data sets

#### Homeostasis and γ-irradiation

Count matrices from Haber et al.(*72*) and Ayyaz et al.(*28*) were downloaded from GEO using GSE92332 and GSE117783, respectively. After preprocessing using the Seurat workflow, Seurat objects were merged and gene expression was averaged by cell type and study using AverageExpression. Next, gene set variation analysis (GSVA)(*64*) was computed using the Gaussian kernel and the output was visualized in a heat map using the ComplexHeatmap package.

#### AIN76A and NWD1 diets: scRNAseq

The intestinal epithelia of mice fed with control (AIN76A) and western-style diets (4 days NWD1, 4 days NWD1 + 4 days AIN recovery) were collected and sent for 10X Genomics scRNAseq analysis (N = 3 mice per group)(*43*). After preprocessing with Seurat and batch correction using the integrated assay workflow, the Seurat clusters were annotated according to canonical markers. CytoTRACE(*45*) was computed on the whole data set using default settings. Next, Paneth cells (N = 991 cells) were taken from the data set, sub-clustered, and visualized by UMAP with a split by the dietary condition. The DSS signature was computed using AddModuleScore and projected on the UMAP embedding. The Paneth sub-cluster that appeared in the NWD1 conditions and associated with the DSS signature was relabeled as ‘Diet response’. Next, marker genes were visualized using ViolinPlot based on the separation of diet response PCs and other PCs. Last, Paneth cells from the diet data set were merged with the Paneth cells from the DSS data set. Average expression was computed based on experimental condition, and GSVA analysis (Gaussian kernel) was performed using the predefined gene sets from the Molecular Signature Database (Hallmark set)(*36, 64*). A filter was applied to select pathways that were both upregulated in DSS and NWD1 condition (ΔGSVA = 0.2). The output was visualized as heat map using the ComplexHeatmap package.

#### AIN76A and NWD1 diets: scATACseq

The intestinal epithelia of mice fed with control (AIN76A) and western-style diets (3 months) were collected and sent for 10X Genomics scATACseq analysis (N = 2 mice per group)(*43*). Processing was done with Signac(*71*), and UMAP embedding was produced base on the 2:30 latent semantic indexing (LSI) components. Unsupervised clustering was performed by a shared nearest neighbor (SNN) modularity optimization based clustering algorithm (algorithm: 1, resolution: 1.2). Cell type predictions scores were obtained by the integration with the AIN76A/NWD1 scRNAseq data set using the FindTransferAnchors and TransferData functions. Two out of twenty-one clusters had high association to the Paneth cell prediction score (cluster 5,20). Paneth cells were defined as part of either cluster 5/20 with the additional condition of a non-zero prediction score for Paneth cells. Next, Paneth cells (N = 945 cells) were taken from the data set, and visualized by UMAP with a split by the dietary condition. Cluster 5 represented cells from both diets while cluster 20 was enriched in NWD1 and therefore relabeled as ‘Diet response’. To identify differentially accessible peaks, FindMarkers was used with logistic regression test. Peaks were filtered based on P < 0.05, visualized with a RegionHeatmap and RIdeogram on the mouse genome, and gene ontology was performed on the closest genes using the EnrichR prackage.

#### Lgr5 ablation with Diphtheria toxin

Count matrices were obtained from GEO (GSE183299) and processing was done with the Seurat workflow(*62*). The revival stem cell (RSC) signature was evaluated using the Addmodulescore function and visualized across experimental groups and cell clusters using the ComplexHeatmap package. A subset was made of intestinal stem cells, and differential expression analysis was performed with FindMarkers. The results were visualized with the EnhancedVolcano package, where marker genes of Paneth cells were highlighted.

#### AOM/DSS-derived tumors

Data from Vega et al.(*32*) was downloaded through GEO (GSE134255) and processed with Seurat(*62*). Differential abundance analysis was performed with MiloR(*73*) workflow using default parameters. The revival stem cell (RSC) signature and the PC Cluster 4 signature were evaluated using the Addmodulescore function and visualized with ViolinPlots.

### CiberSortx analysis of AOM/DSS adenomas

CiberSortx(*34*) was leveraged to predict the fractions of various cell types in tumors arising in AOM/DSS context. To this aim, count data from AOM/DSS-derived adenomas from Chen et al.(*31*) (GSE178145) were converted to counts per million with Edger. The single cell count data from Paneth cell upon genetic targeting were provided as reference and converted to a signature matrix in CiberSortx using default parameters. For cell type imputation, CiberSortx was applied in relative mode using 100 permutations.

## Data availability

All data relevant to this study are made available. Transcriptomic sequencing data has been deposited in the Gene Expression Omnibus (GEO), and is available using the following identifiers: GSE221819 (bulk RNA sequencing of murine tumors); GSE221820 (single cell RNA sequencing of genetically targeted Paneth cells); GSE221818 (bulk RNA sequencing of sorted Paneth cells treated with control and western-style diet). Additional data sets referenced in this study are publicly available in the Synapse and GEO repositories. Previously published scRNAseq studies were employed relative to the mouse small intestine in homeostasis (Haber et al.(*72*), GSE92332), upon irradiation (Ayyaz et al.(*28*), GSE117783), *Lgr5* ablation (Singh et al.(*30*) GSE183299), and upon feeding with western-style diet (Choi et al. (*43*), scRNAseq: GSE188577, scATACseq: GSE228006). Mouse colonic scRNAseq data from AOM/DSS tumors was obtained from Vega et al. (*32*) (GSE134255) and bulk RNAseq was from Chen et al.(*31*) (GSE178145). Human colorectal cancer Bulk RNAseq and scRNAseq studies were retrieved from Guinney et al.(*38*) (syn2623706), Rajamäki et al.(*35*) and Lee et al.(*37*) (KUL cohort: GSE144735; SMC cohort: GSE132465), respectively. Human scRNAseq data from Ulcerative Colitis patients (Smillie et al.(*51*)) was downloaded from the Broad DUOS platform after a DTA agreement. Single cell ATAC data from human colon was obtained from Becker et al.(*50*) (GSE201349).

## Acknowledgements

We thank E. Bindels and R. Hoogenboezem for their support to generate the single cell sequencing data, J.D. Windster for help with in situ hybridizations, W.S. van de Geer for analysis support of bulk sequencing data; W. Szymanska, E. Middendorp Guerra and E. de Vet for help with the IHC and scRNAseq analyses; V.H.A.E. Verhagen for discussions on XGBoost machine learning. This study was financially supported by the Dutch Cancer Society (KWF; project no. 11407), the Wereld Kanker Onderzoek Fonds and World Cancer Research Funds (WKOF/WCRF; projects no. 2014-1181 and IIG_FULL_2022_015), and in part by the National Cancer Institute (NCI; project no. R01CA214625, R01CA229216, and P30-013330), the National Institute on Aging (NIA; project no. P30 AG038072, 5T32AG023475-20), Academy of Finland (Finnish Center of Excellence Program 2018–2025 312041, Academy Professor grant no. 319083 and 320149), iCAN Digital Precision Cancer Medicine Flagship (320185), Cancer Society of Finland, Sigrid Jusélius Foundation (220002), and the Jane and Aatos Erkko Foundation.

## Author contributions

M.P.V., R.J., M.S. and A.S. designed and performed experiments. M.P.V., J.C., N.V., L.A.A. and L.H.A. generated and analyzed sequencing data sets. K.R., P.P, S.S, B.v.d.S. and T.P.P.v.d.B. performed immunohistochemistry. D.S. and M.D. were responsible for pathological examination and IBD patient selection. M.P.V. and R.F. wrote the manuscript. R.F. supervised the study and was responsible for concept and design of the study.

## Supplementary Figure Legends

**Supplementary Figure 1.**
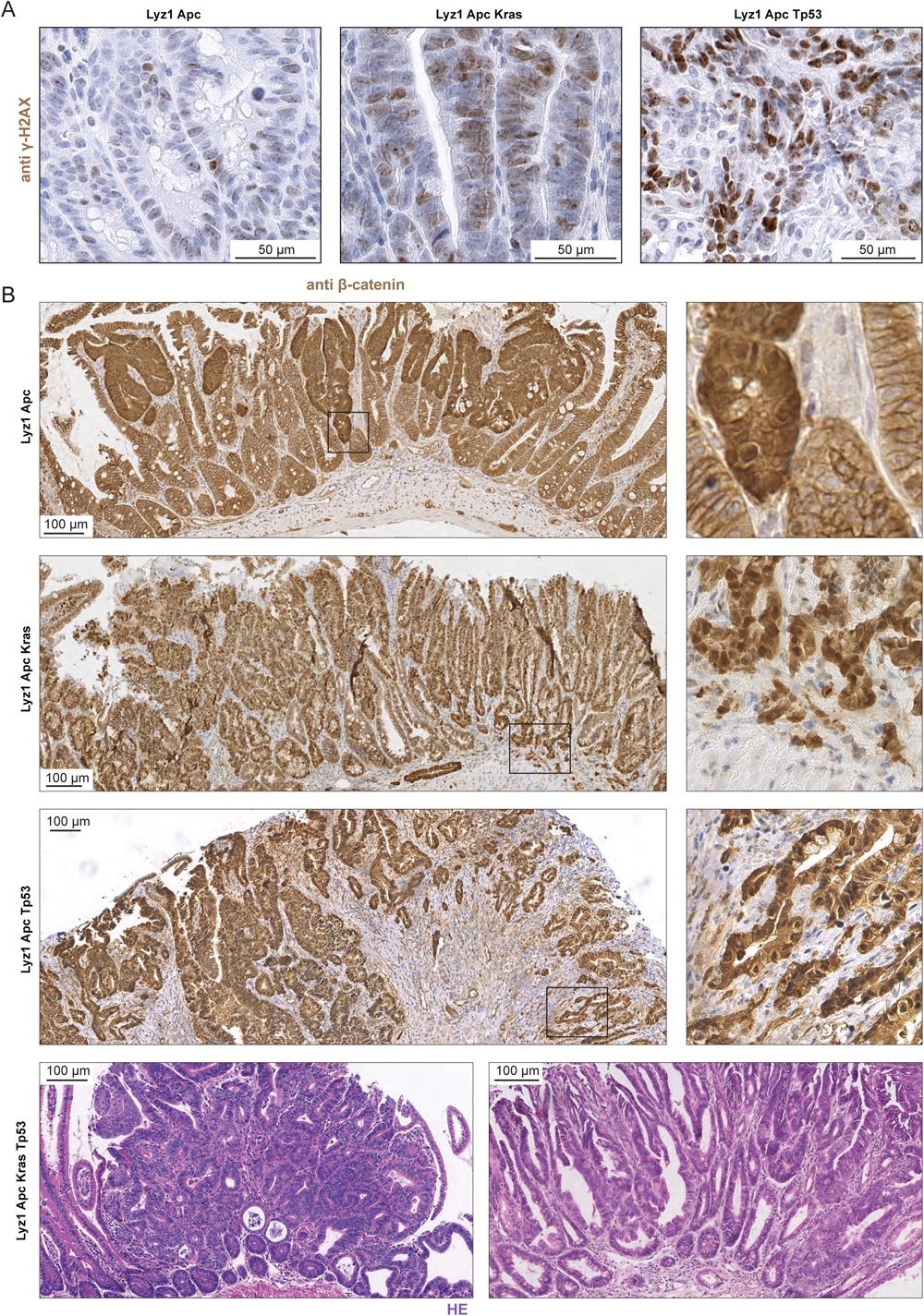
Immuno-histologic analysis of Paneth-derived tumors. **a.** Representative IHC images of Paneth-derived tumors from different genetic backgrounds, stained with phospho-histone H2A.X (Ser 139). **b.** Top three panels: IHC analysis of Paneth-derived intestinal tumors with different genetic backgrounds (as indicated), stained for β-catenin. Bottom: hematoxylin and eosin histological stains of Paneth-derived tumors from *Lyz1*/*Apc*/*Kras*/*Tp53* mice.

**Supplementary Figure 2.**
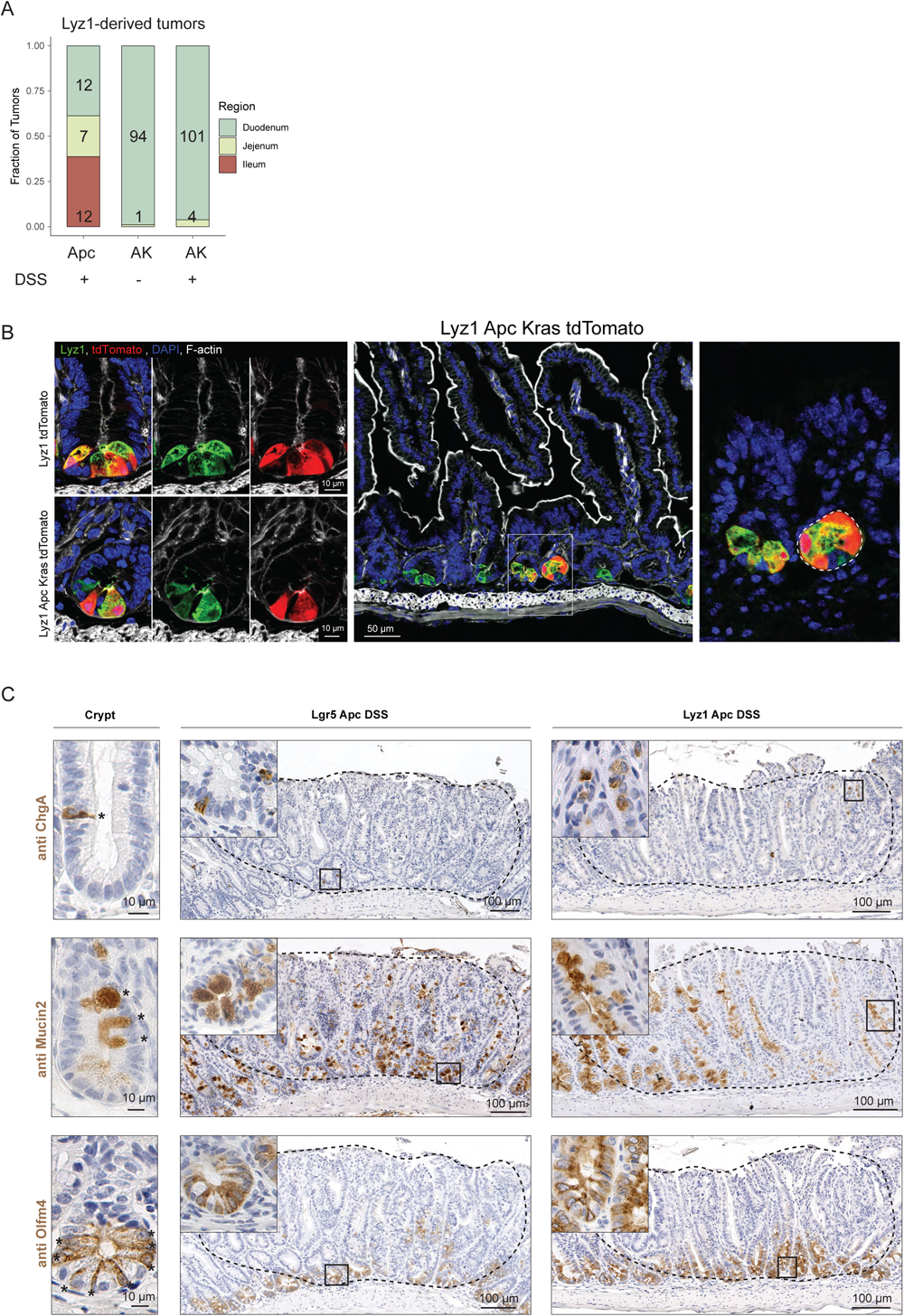
Characterization of PC-derived small intestinal adenomas. **a.** Stacked bar plot displaying the localization and distribution of PC-derived adenomas across the small intestine. **b**. Immunofluorescence imaging of intestinal crypts from *Lyz1*/tdTomato and *Lyz1*-*AK*/tdTomato mice. **c**. IHC analysis of the lineage-specific ChgA, Muc2, and Olfm4 markers in PC-and *Lgr5*-derived adenomas.

**Supplementary Figure 3.**
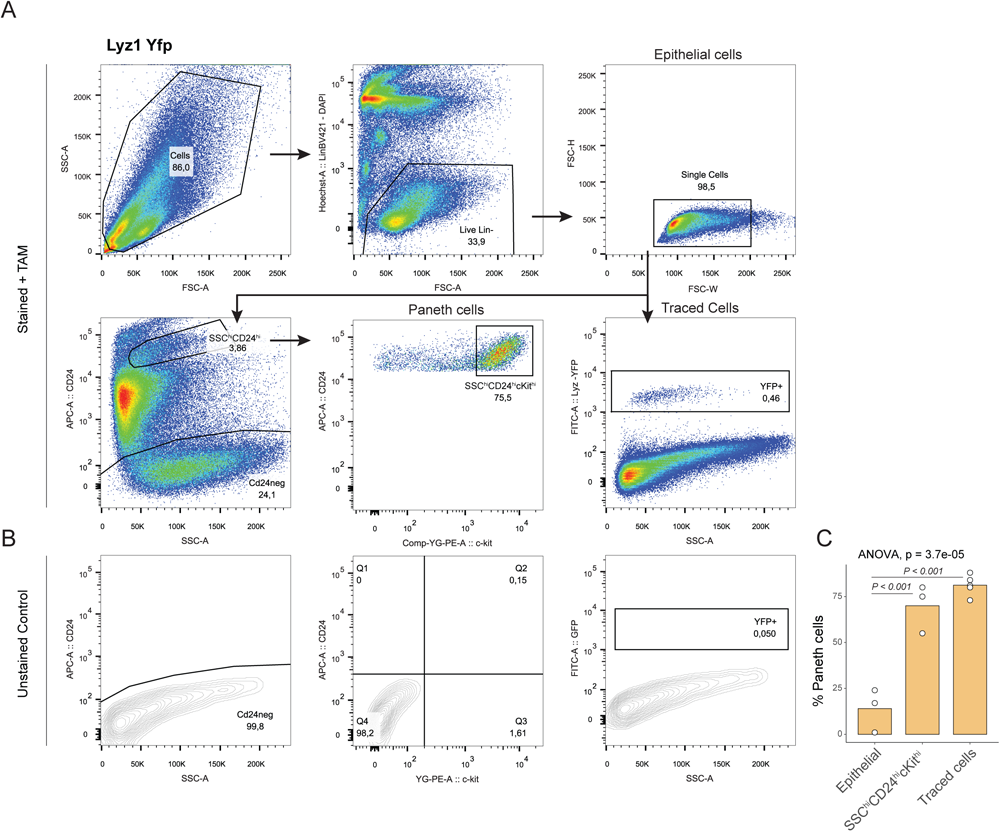
FACS gating strategy for Paneth cells enrichment and isolation. **a.** FACS gating strategy established for the isolation of live, single, epithelial cells (top). Subsequently, Paneth cells were enriched using the SSC^hi^CD24^hi^ gate and further purified based on high cKit levels (SSC^hi^CD24^hi^cKit^hi^). Alternatively, traced cells were sorted based on positive expression for Yfp or td-Tomato when the appropriate reporter mice were employed. **b.** Negative control employed to establish reference levels based on unstained samples. **c.** Bar plot of the percentage of Paneth cells obtained by the distinct gating strategies. P values depict results of one-way ANOVA and Tukey tests.

**Supplementary Figure 4.**
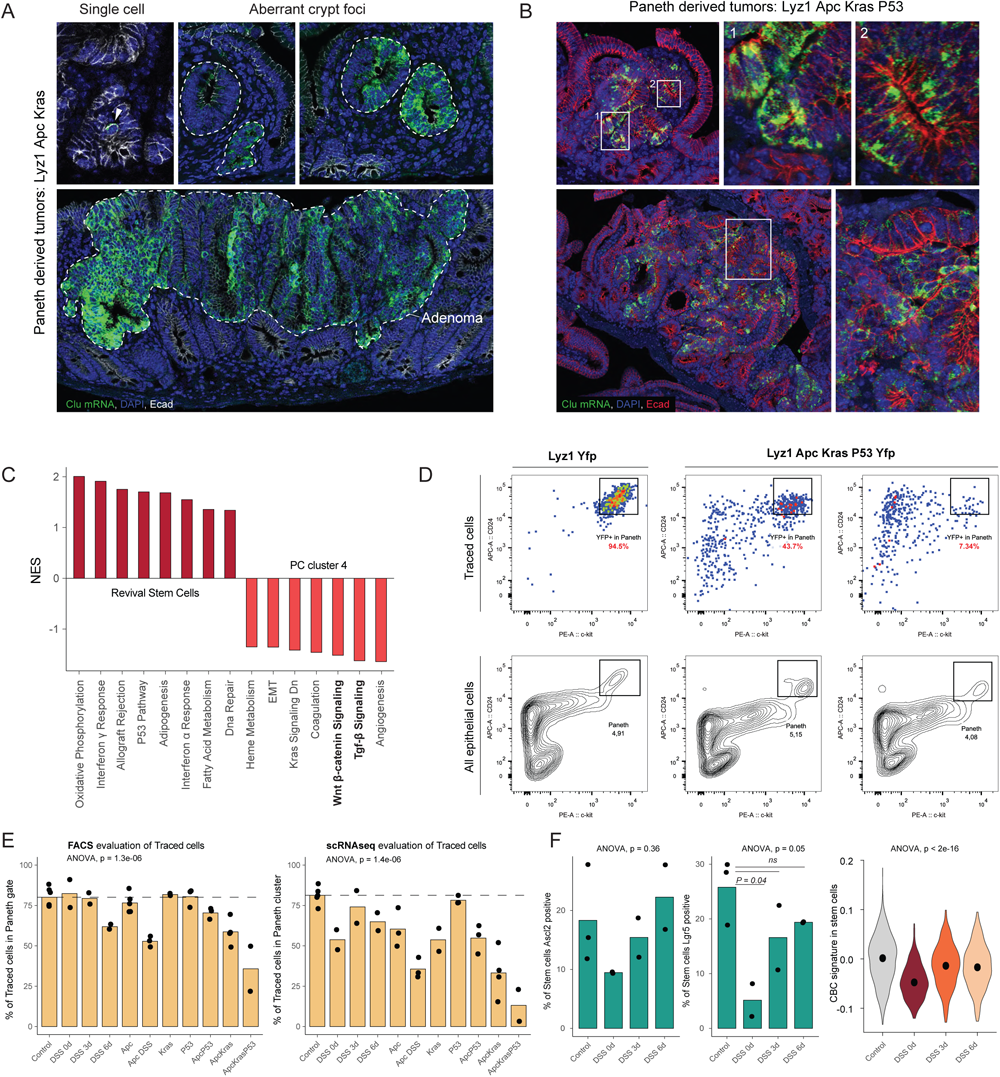
Paneth cells dedifferentiate upon DSS-driven inflammation. **a-b.** In situ hybridization (ISH) analysis of Clu mRNA (green) expression at various stages during PC-derived tumorigenesis in *Lyz1*-*Apc*/*Kras* and (**b**) *Lyz1*-*Apc*/*Kras*/*Tp53* mice. **c.** Bar plot showing filtered pathways (Pval < 0.05, abs NES > 0.5) of the gene set enrichment analysis comparing RSCs with PC Cluster 4. **d.** FACS analysis of lineage-traced (Yfp+) cells in wild type (*Lyz1*-Yfp) and AKP-mutant (*Lyz1*-*Apc*/*Kras*/*Tp53*) mice. The latter clearly localize outside of the CD24^hi^cKit^hi^ gate where wild type PCs reside. **e.** Bar plot relative to the percentages of lineage-traced cells evaluated by FACS using the CD24^hi^cKit^hi^ PC-specific gate (left), or by scRNAseq analysis (% within Paneth cluster; right). P values denote significance of one-way ANOVA. **f.** Left: Bar plots showing the average percentage of *Ascl2*^+^ and *Lgr5*^+^ stem cells in control and DSS-treated animals. Right: Violin plots showing a decrease in the crypt base columnar (CBC) signature upon DSS treatment. P values denote significance of one-way ANOVA and Tukey test for group comparisons.

**Supplementary Figure 5.**
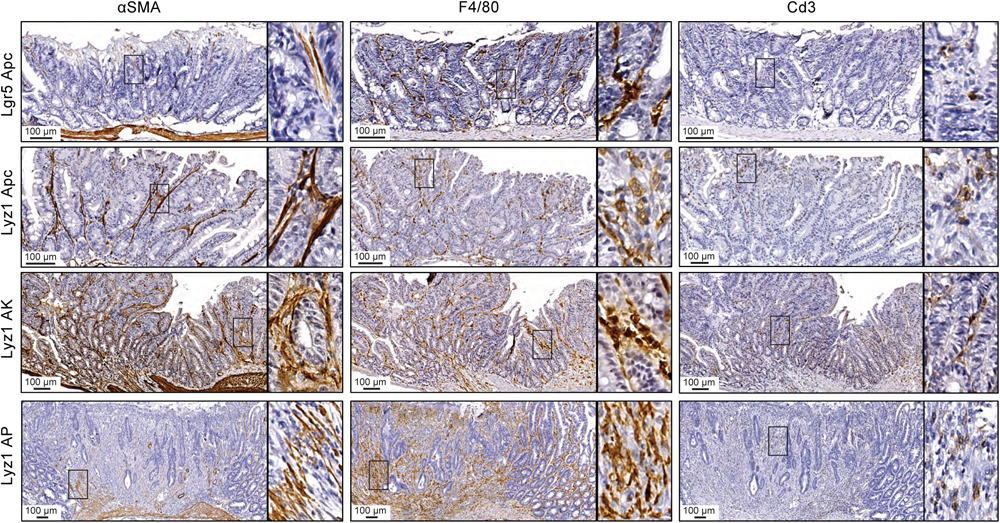
Distinct TME features in CBC-and PC-derived small intestinal adenomas. IHC analysis of *Lgr5*^+^- and *Lyz*-derived adenomas with the tumor microenvironment markers αSMA (fibroblasts), F4/80 (macrophages), and Cd3 (T cells).

**Supplementary Figure 6.**
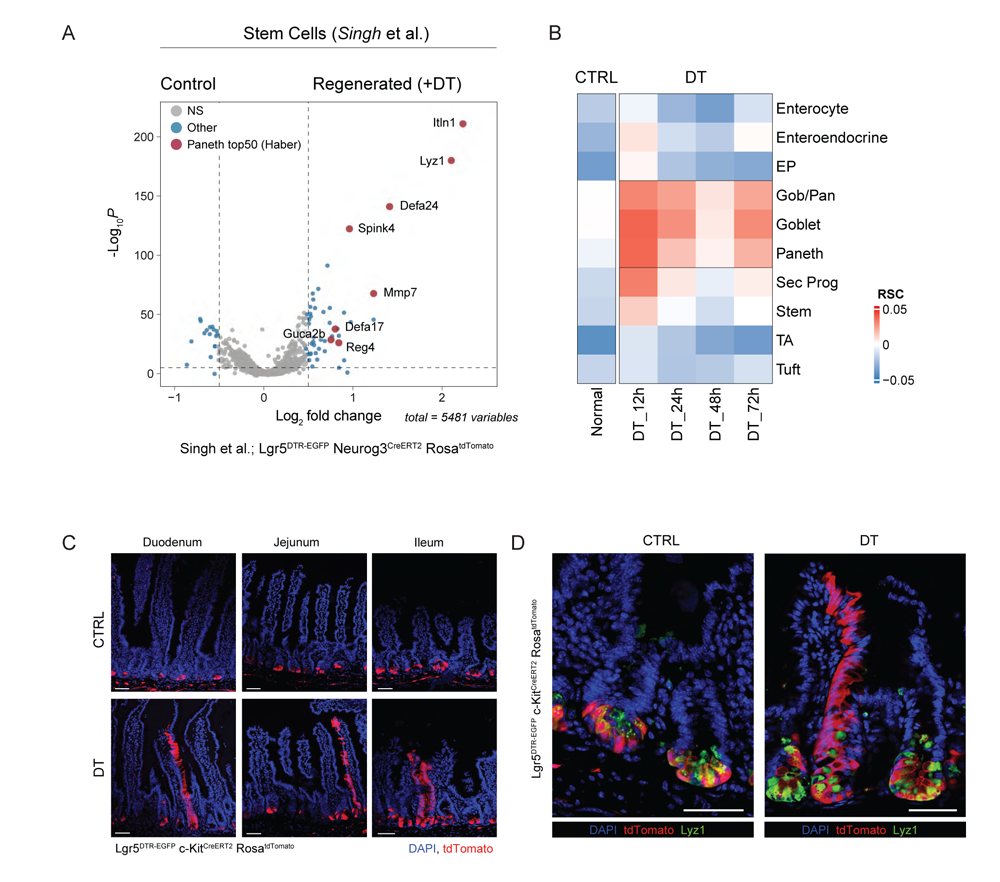
*Lgr5*^+^ CBCs ablation triggers dedifferentiation of Paneth and goblet secretory lineages. **a.** Differential expression analysis of intestinal stem cells in homeostasis and during regeneration following *Lgr5*^+^ CBCs ablation. scRNAseq are from Singh et al.(*30*). Several among the top 50 differentially expressed genes in the regenerated ISCs are Paneth-specific markers (red). **b**. Heatmap showing transcriptional activity of the RSC program upon *Lgr5*^+^ CBCs ablation. **c-d**. Lineage tracing of DT-treated *Lgr5*^DTR-EGFP^;*c-Kit*^CreERT2^; Rosa^tdTomato^ mice across different intestinal sections (c.) and double stained with Lyz1/tdTomato (d.). Scale bar = 50 μM, 2 mice per group were analyzed.

**Supplementary Figure 7.**
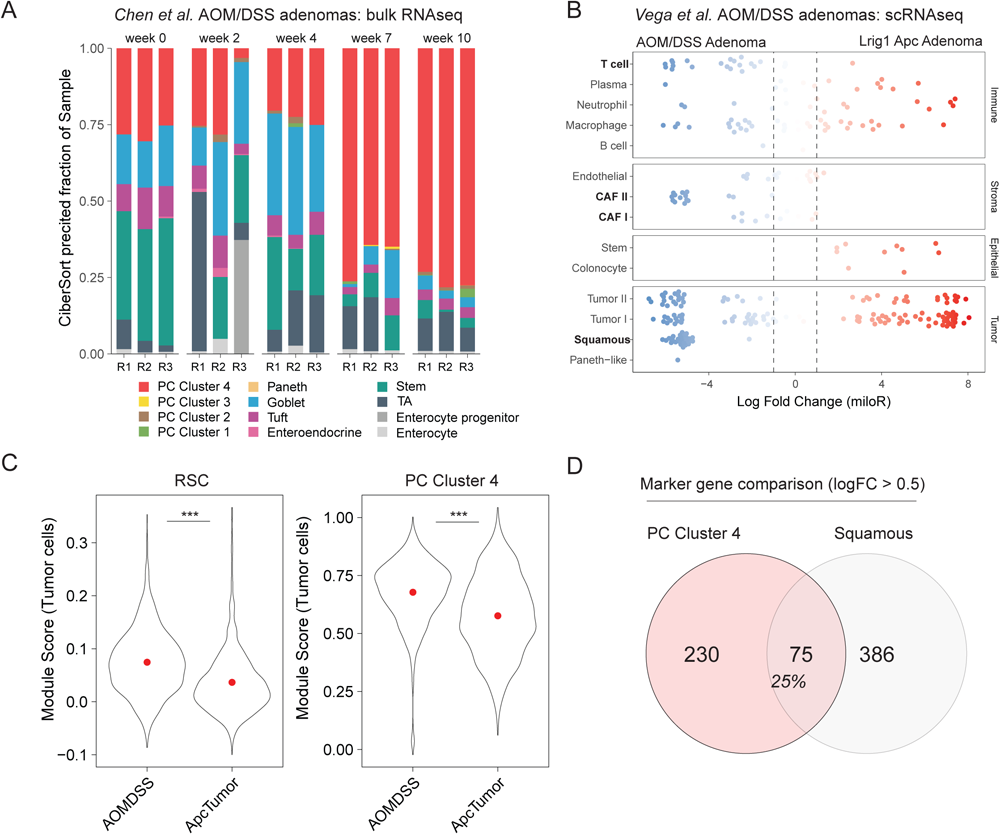
AOM/DSS-derived adenomas share similarities with PC-derived tumors. **a.** Stacked bar plot of the cell type fractions in AOM/DSS adenomas as predicted by Cibersort analysis(*34*) (see Methods). Bulk RNAseq data are from Chen et al.(*31*). **b**. Differential cell type abundance (MiloR(*73*); see Methods) between AOM/DSS and *Lrig1*-*Apc* adenomas. scRNAseq data are from Vega et al.(*32*). **c**. Comparison of RSC and PC Cluster 4 module score in tumor cells from AOM/DSS and *Lrig1*-*Apc* adenomas. **d**. Venn diagram indicating the overlap between marker genes from PC Cluster 4 and the AOM/DSS-specific squamous tumor cell subpopulation.

**Supplementary Figure 8.**
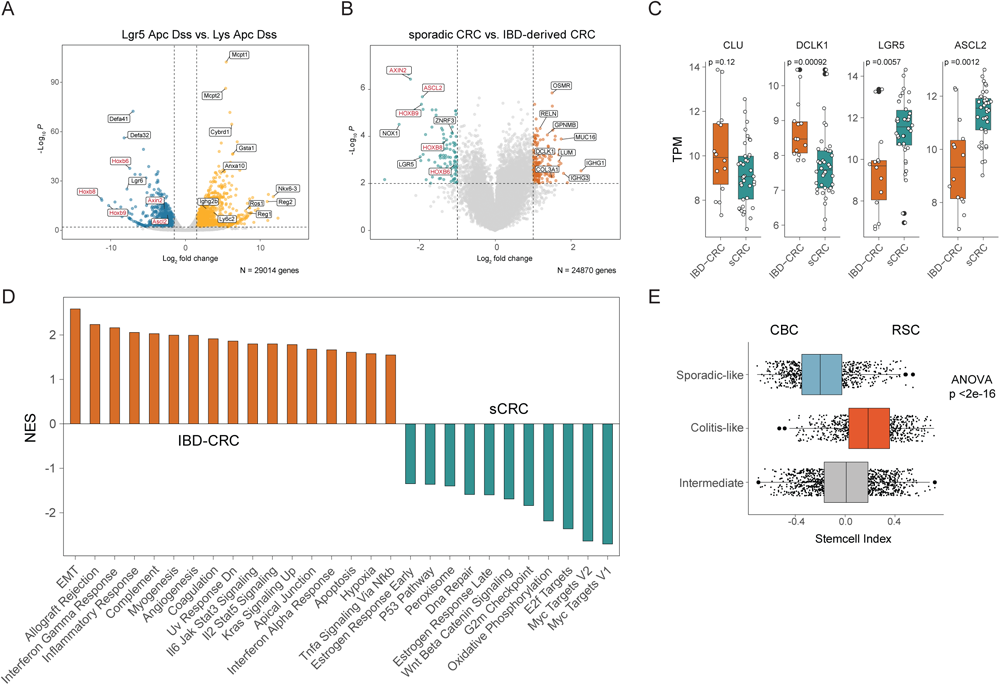
Differential gene expression analysis between IBD colon cancers and murine PC-derived small intestinal adenomas. **a.** Volcano plot showing differentially expressed (Pval < 0.01, FC cutoff 1.5) genes between *Lgr5*-and Paneth-derived intestinal tumors. **b**. Volcano plot highlighting differentially expressed (Pval < 0.01, FC cutoff 1) genes between sporadic-and IBD-CRCs. **c**. Box plots relative to differentially expressed CBC (Lgr5-derived) and RSC (PC-derived) marker genes between sporadic-and IBD-CRCs. P values denote results of the t test. **d**. Bar plot relative to the gene set enrichment analysis between sCRC and IBD-CRC (abs NES > 0.5, Pval < 0.05). **e**. Box plots relative to stem cell index values across the colitis, sporadic, and intermediate group of colon cancers. P value depicts result of one-way ANOVA.

**Supplementary Figure 9.**
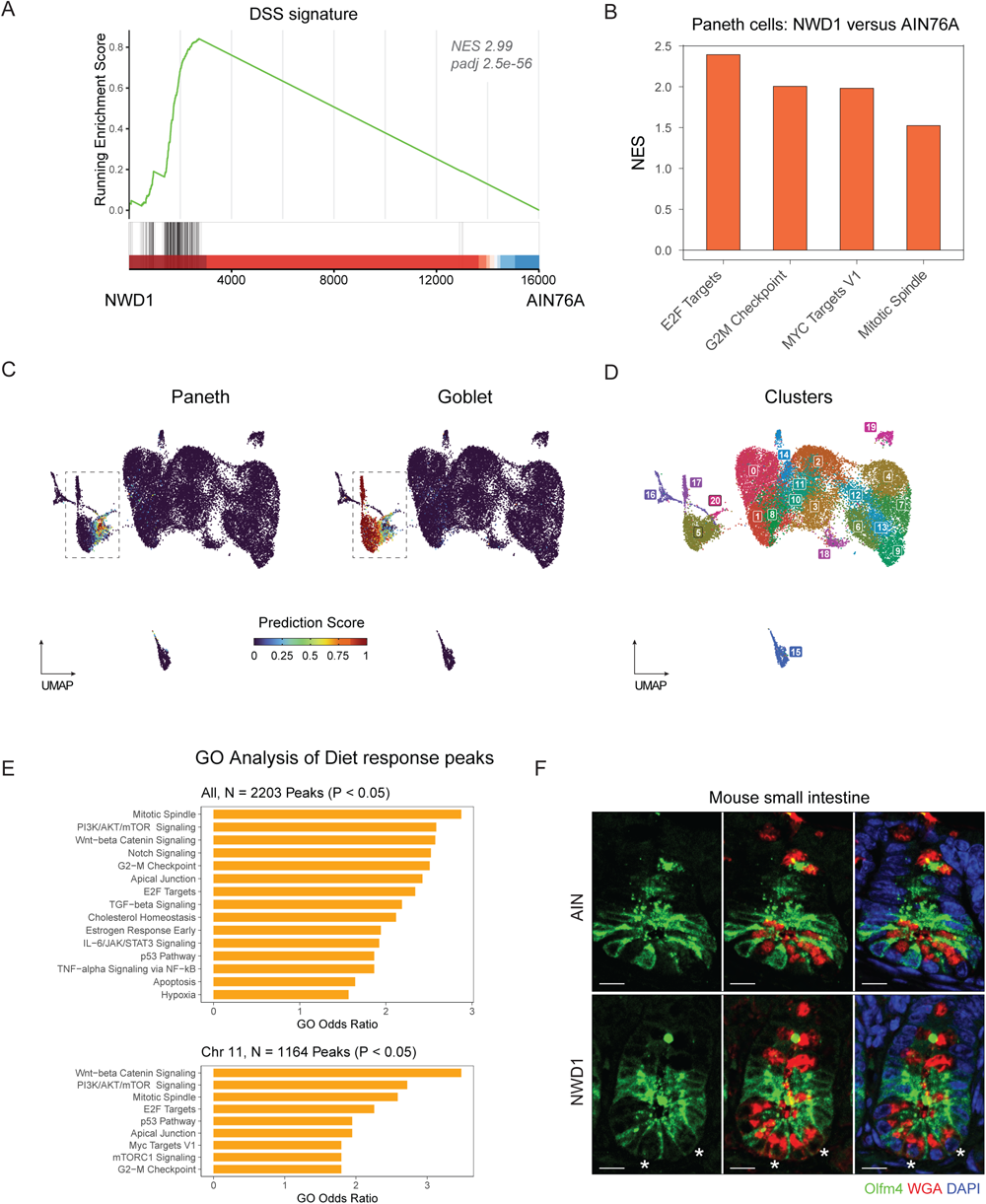
The NWD1 western-style diet triggers the dedifferentiation of Paneth cells. **a.** GSEA plot of Paneth cells from NWD1-vs. AIN76A-fed mice. The DSS signature significantly associates with the NWD1 diet (NES 2.99, Padj 2.5e-56). **b**. Bar plot showing pathways elevated (P < 0.05, NES > 0.5) in Paneth cells from NWD1-when compared to AIN76A-fed animals. **c**. Prediction scores for Paneth (left) and goblet (right) cells on UMAP embedding from the scATAC data (Choi et al.(*43*)). **d**. Unsupervised clustering reveals two distinct clusters (c5, c20) that include Paneth cells. **e**. Bar plot listing gene ontology results based on differential peaks from the diet responsive Paneth cells. **f**. Immunofluorescence imaging of WGA and Olf4m showing intestinal crypts from mice fed with AIN/NWD1 diets. Asterisks earmark double positive cells. Scale bar indicates 10 µm.

**Supplementary Figure 10.**
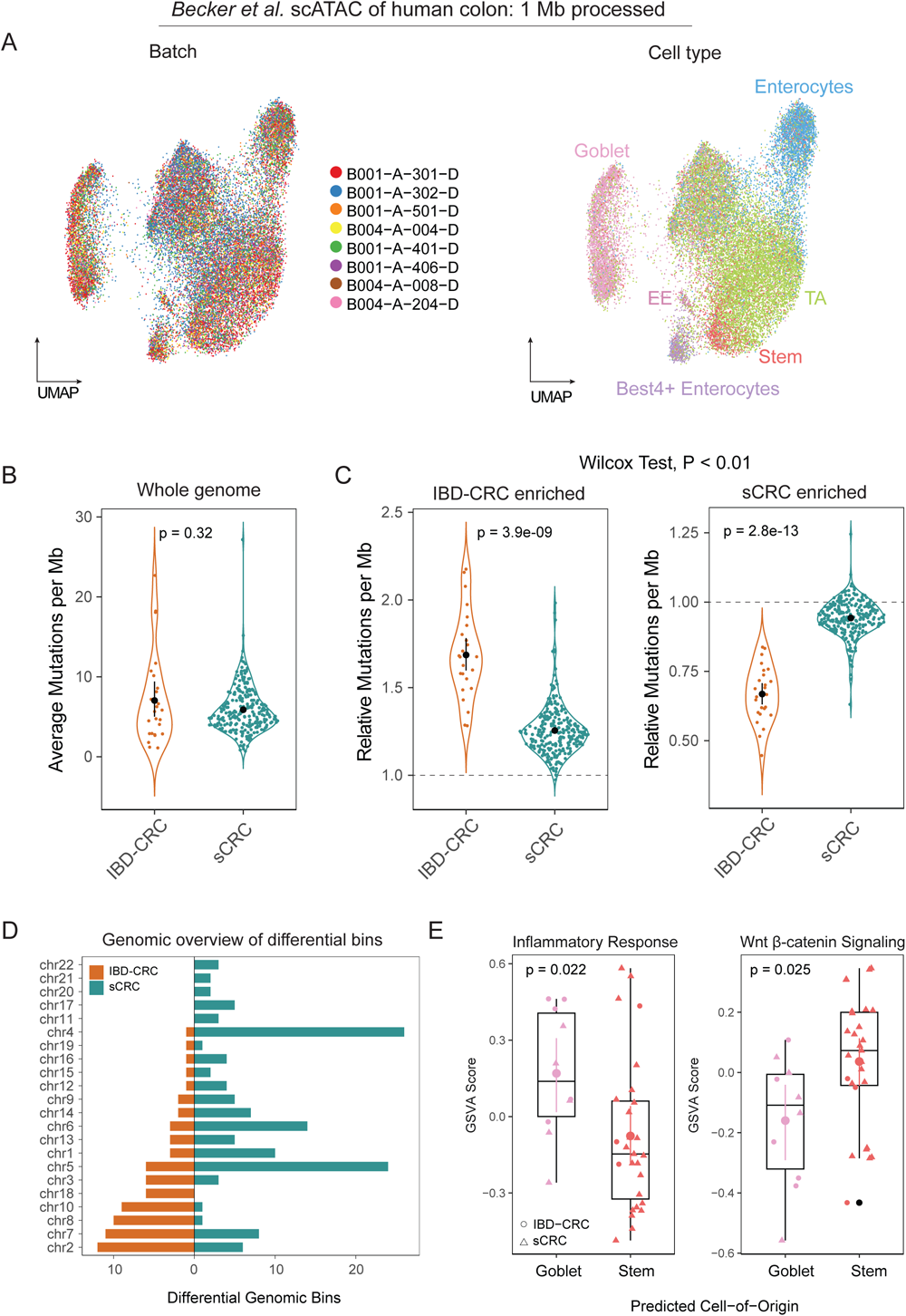
Epigenetic profiles of normal colonic lineages and mutational patterns of IBD-related and sporadic colon cancers. **a.** UMAP plot displaying 1 Mb processed scATAC data of the human colon (Becker et al.(*50*)). **b**. Average number of passenger mutations in IBD-related and sporadic colon cancers(*35*). **c**. Relative mutation frequency in IBD-enriched (N = 78) and sCRC-enriched (N = 136) bins based on Wilcox test (p<0.01). **d**. Barplot showing the genomic overview of differential bins across chromosomes. **e**. Boxplots relative to the gene set variation analysis (GSVA) scores of tumors stratified according to their predicted cell of origin. P values denote results of t tests.

